# Epithelial Plasticity and Innate Immune Activation Promote Lung Tissue Remodeling following Respiratory Viral Infection

**DOI:** 10.1101/2021.09.22.461381

**Authors:** Andrew K. Beppu, Juanjuan Zhao, Changfu Yao, Gianni Carraro, Edo Israely, Anna Lucia Coelho, Cory M Hogaboam, William C. Parks, Jay K. Kolls, Barry R. Stripp

## Abstract

Epithelial plasticity has been suggested in lungs of mice following genetic depletion of stem cells but is of unknown physiological relevance. Viral infection and chronic lung disease share similar pathological features of stem cell loss in alveoli, basal cell (BC) hyperplasia in small airways, and innate immune activation, that contribute to epithelial remodeling and loss of lung function. We show that a novel subset of distal airway secretory cells, intralobar serous (IS) cells, are activated to assume BC fates following influenza virus infection. Injury-induced hyperplastic BC (hBC) differ from pre-existing BC by high expression of IL-22Ra1 and undergo IL-22-dependent expansion for colonization of injured alveoli. Resolution of virus-elicited inflammation resulted in BC>IS re-differentiation in repopulated alveoli, and increased local expression of antimicrobial factors, but failed to replace normal alveolar epithelium. Epithelial plasticity therefore protects against mortality from acute respiratory viral infection but results in distal lung remodeling and loss of lung function.

## Introduction

Stem cell plasticity contributes to tissue regeneration and includes a range of processes such as lineage conversion and de-differentiation of specialized progenitor cells (Blanpain and Fuchs, 2014). Although epithelial cell plasticity in lung repair has been inferred from lineage tracing and targeted cell ablation studies (Tata et al., 2013; Tata and Rajagopal, 2017), the physiological relevance in lung disease is unknown. The epithelial linings of airways and alveoli are maintained by distinct regional facultative stem/progenitor cells whose progeny include both self-renewing and differentiating subsets (Hogan et al., 2014; Rackley and Stripp, 2012). Regional differences in the fate of differentiating progeny allow for maintenance of locally specialized epithelial functions, such as mucociliary clearance and host defense in the conducting airways and gas exchange in the distal respiratory units. Restriction of progenitor cells and their differentiating progeny to distinct anatomic zones during homeostatic tissue maintenance are necessary for functional integration of these compartments along the proximodistal axis and preservation of normal physiological lung function. However, acute lung injury and chronic lung disease disrupt normal progenitor cell compartmentalization leading to aberrant tissue remodeling and declining lung function. Such is the case following infections by respiratory viruses such as H1N1 influenza A or SARS-CoV2, and in chronic lung diseases such as idiopathic pulmonary fibrosis (IPF). In such conditions, basal cell (BC) hyperplasia in airways leads to the recruitment and colonization of these cells into injured alveolar areas, and this proximalization of distal lung tissue contributes to potentially life-threatening loss of alveolar diffusion capacity (Seibold et al., 2013; Vaughan et al., 2015; Xu et al., 2016; Zuo et al., 2015). However, the identity of epithelial progenitor cells that contribute to proximalization of distal lung tissue and the mechanisms that regulate their fate during tissue remodeling remain poorly defined.

Mirroring what occurs in injured or infected human lungs, acute lung injury in mice infected with a mouse-adapted Puerto Rico 8 (PR8) variant of H1N1 influenza virus is accompanied by BC expansion in airways that ultimately replaces the injured epithelium of the alveolar gas-exchange region (Kumar et al., 2011; Vaughan et al., 2015; Zuo et al., 2015). Even though basal and club cells serve as stem cells for maintenance of the pseudostratified epithelium of proximal airways and cuboidal epithelium of distal airways, respectively, lineage tracing studies suggest that hyperplastic BC appearing in distal lung tissue of PR8-infected mice are derived from neither of these canonical stem cell populations (Ray et al., 2016; Vaughan et al., 2015; Zuo et al., 2015). Instead, injury-induced hyperplastic BC (hereon referred to as hBC) derive from alternate small airway progenitors that can be lineage traced based upon expression of either Sox2 or p63 transcription factors (Ray et al., 2016; Xi et al., 2017; Yang et al., 2018). Similarly, in proximal airways, α-smooth muscle actin-expressing myoepithelial cells of submucosal glands (SMG) or SSEA4-expressing secretory cell progeny of BC can replenish basal stem cells following injury (Tata et al., 2018; Tata et al., 2013).

However, the molecular and functional relationship between epithelial progenitors of SMG or upper airway surface epithelium that can replace local basal stem cells, versus those that yield expanding BC in small airways following PR8 infection, remain to be established.

Although much is known of mechanisms that regulate the renewal and fate of BC within pseudostratified airways, mechanisms regulating hyperplastic BC appearing in alveolar epithelium of PR8-infected mice are poorly defined. Evidence of roles for local hypoxia (Xi et al., 2017) and the altered inflammatory milieu (Katsura et al., 2019; Pociask et al., 2013; Tavares et al., 2017) that accompanies PR8 infection suggest potential roles for innate immune activation as a regulator of BC fate and epithelial remodeling. Epithelial-immune crosstalk serves as a critical regulator of progenitor cell function in multiple organs including the lung, gut, and skin (Aparicio-Domingo et al., 2015; Barrow et al., 2018; Boniface et al., 2005; Kempski et al., 2017; Lindemans et al., 2015). Here, we show that the unique origins and molecular phenotype of hBC elicited by PR8 infection allows for their dynamic response to the activated innate immune system. Interleukin 22 derived from locally activated γδT cells promotes self-renewal and hyperplasia of BC within alveolar epithelium that ultimately assume serous cell fates in previously injured regions during resolution of the PR8-elicited inflammatory response. This remodeling response to respiratory viral infection allows for efficient replacement of exposed basement membrane in the injured alveolar epithelium and local production of an antibacterial secretome to protect against secondary bacterial infection.

## Results

### Activation and expansion of intralobular serous (IS) cells precedes BC hyperplasia following influenza-induced acute lung injury

BC hyperplasia occurs in airways of patients with chronic lung disease and in lung tissue of those succumbing to acute respiratory viral infections such as H1N1 influenza virus or COVID-19 (Davis and Wypych, 2021; Fang et al., 2020; Rock et al., 2010; Vaughan et al., 2015; Zuo et al., 2015). Based on lineage tracing studies, the majority of cells in mouse lungs that acquire BC fates in response to influenza virus-induced acute lung injury have been proposed to arise from immature P63^+^Krt5^-^ basal progenitors (Xi et al., 2017; Yang et al., 2018). However, a significant fraction of hBC seen during injury failed to retain a P63 lineage tag suggesting the presence of other contributing non-basal progenitors (Fernanda de Mello Costa et al., 2020). We sought to interrogate the existence of non-canonical progenitors by assessing dynamic changes in epithelial cell populations in response to influenza-induced acute lung injury. We generated a comprehensive profile of single cell transcriptomes for epithelial cells isolated from trachea, extrapulmonary bronchus, intralobar airways and alveolar regions of control C57Bl/6 mice (Day 0) and of mice infected and recovering from exposure to the PR8 strain of H1N1 influenza virus (Day 3-240; Fig. 1A, B). Results displayed as uniform manifold approximation and projection (UMAP) two-dimensional plots reveal eight major cell clusters that were categorized according to known lung epithelial cell types based upon their unique gene expression signatures (Fig. 1B, C; Supp. Fig. 1A). Even though each of the major cell types were observed among epithelial cells sampled from all control and post-exposure time points, their relative proportions within total sampled epithelial cells showed significant variability (Supp. Fig. 1B). Basal cells were only infrequently observed among epithelial cells sampled from lobes of control mice but showed progressive increases in representation at time points evaluated following PR8 infection (Fig. 1D, E). Serous cells were the only other rare epithelial cell type in lung lobes of naïve mice whose abundance increased following PR8 infection (Fig. 1D, E). However, the increase in representation of serous cells preceded increases in BC representation among epithelial cells sampled across the time course (Fig. 1E).

**Fig. 1.**
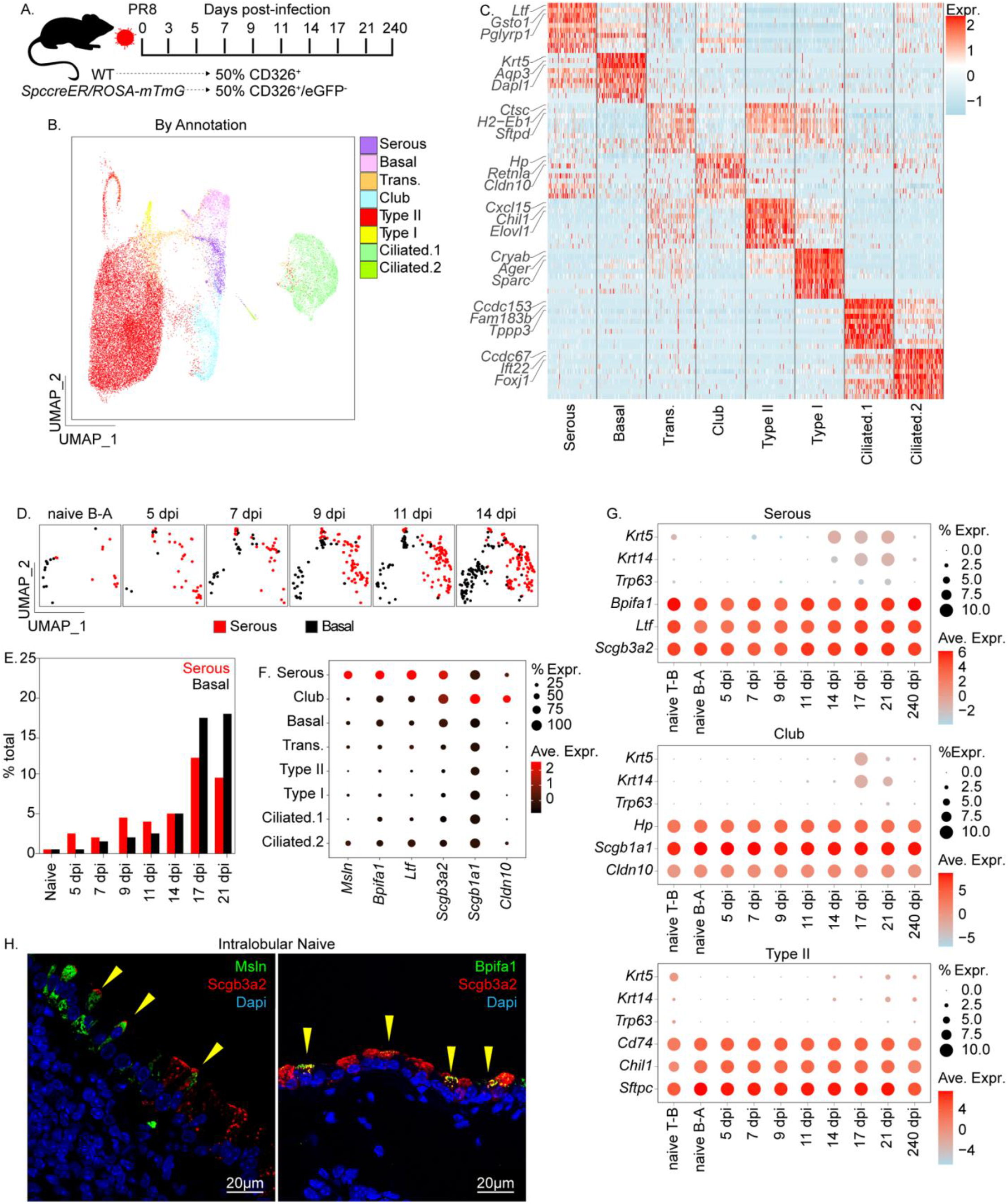
ScRNAseq to reveal the molecular phenotype of lung epithelial cells during the course of recovery following PR8 influenza virus infection. (A) Experimental design. Mouse lung homogenates were collected from 8–12-week-old C57/Bl6 and *Sftpc-CreER*/*ROSA-mTmG* mice (n = 5 per group). Lung tissue was collected at indicated time points and either total epithelial cells (C57/Bl6 mice;CD31^-^CD45^-^CD326^+^) or AT2 depleted epithelial cells (*Sftpc-CreER*/*ROSA-mTmG* mice; CD31^-^CD45^-^CD326^+^eGFP^+^) enriched by FACS. (B) UMAP plot of combined scRNAseq data. Unsupervised clustering was used to distinguish distinct cell phenotypes which were assigned to known epithelial cell types based upon gene signatures. Pie chart shows fractional representation of each cell types within the entire data set. (C) Heatmap showing unique molecular profiles between cell types. Top 3 genes for each cell category are annotated to the left. Full gene lists for each cell category are provided in Supp. Fig. 1A. (D) UMAP plot tracking relative changes in basal and serous cell populations for indicated samples. B-A = bronchioalveolar. (E) Representation of serous (red bars) and basal (black bars) cell types as a function of percent total sampled cells at each recovery time point. Analysis was performed on AT2 depleted samples (ie using the FACs enrichment strategy CD31^-^CD45^-^CD326^+^eGFP^+^ from *Sftpc-CreER*/*ROSA-mTmG* mice). dpi = days post infection. (F) Dot plot comparing expression of selected club and serous cell-specific genes between cell types. (G) Assessment of BC gene signature during recovery from PR8-induced injury. T-B = tracheal bronchial. (H) Representative immunofluorescence colocalization of either Msln or Bpifa1 (green) with Scgb3a2 (red) within conducting airway epithelium.

Because serous cells show increased representation in single cell data sets at earlier stages in the response to PR8 infection, we speculated that serous cells represent the progenitor cell type accounting for hBC. To explore further the molecular phenotype of serous cells that appear early in the airway response to PR8 infection, we interrogated our scRNAseq data to identify differentially expressed genes (DEGs) that discriminate serous cells from other epithelial cell types in conducting airways. Expanding populations of serous cells observed during recovery following PR8 infection were unique among distal lung epithelial cell types in their expression of genes involved in anti-microbial host defense, such as *Ifitm1, Ifitm3, Bpifa1*, and *Ltf* (Supp. Fig. 1C). Despite many similarities in gene expression signatures between club and serous cells, unsupervised clustering of these epithelial populations was unaffected by regression of cell cycle genes (Supp. Fig. 1D, E, F). We found that serous cells express high levels of *Scgb3a2*, as do club cells, but that unlike club cells they lack expression of *Scgb1a1*. Of note, antimicrobial factors, such as *Bpifa1* and *Ltf*, whose expression defines serous cells of proximal airways and submucosal glands (Tata et al., 2018), were significantly elevated in serous cells compared to club cells of the PR8 injured lung (Fig. 1F; Supp. Fig. 2A). Furthermore, serous cells of the PR8-exposed lung include a subset that harbor transcripts for BC marker genes including *Trp63, Krt14* and *Krt5*, present at variable levels among all post-exposure recovery time points but absent in airways of naïve mice (Fig. 1G). Immunofluorescence staining revealed that a small fraction of intralobular Scgb3a2-immunoreactive cells were also positive for Msln and Bpifa1, indicating the presence of serous populations within the lower respiratory tract, herein called intralobular serous (IS) cells (Fig. 1H). Expansion of IS cells coupled with acquisition of a transcriptome that shares similarities with nascent airway BC during recovery from PR8 exposure, led us to speculate that rare airway serous cells represent the progenitor cell-of-origin for hBC observed in airways and alveolar epithelium of infected mice.

### Transcriptome analysis predicts IS>basal plasticity during recovery from PR8 infection

To further examine the potential for IS cells to serve as progenitors for expansion of hBC, we used gene set enrichment to identify DEGs among these epithelial cell clusters at different times post-PR8 infection. We observed a significant induction of gene sets associated with cell cycle at 5- and 7-days post infection coinciding with the appearance of proliferating conducting airway epithelium observed *in vivo* (Fig. 2A). However, we observed enrichment of cell-cycle gene-sets at these time points only in serous and BC with little to no enrichment of these gene sets in other major cell types of conducting airways (Fig. 2A). We used our scRNAseq dataset to interrogate lineage relationships between BC, IS cells, and other airway secretory cell populations. Transcript splicing was assessed by Velocyto (La Manno et al., 2018) to predict differentiation trajectories among IS cells, club cells, and hBC during early (naïve-day 14) and late (day 14-day 21) responses to PR8 infection (Fig. 2B, C). Computational analysis of cell fate dynamics predicted a process that IS cells de-differentiated into BC at early time points post-PR8 infection (Fig. 2C). In contrast, trajectory analysis predicted the transition of BC back to IS cells at late recovery time points (Fig. 2C).

**Fig. 2.**
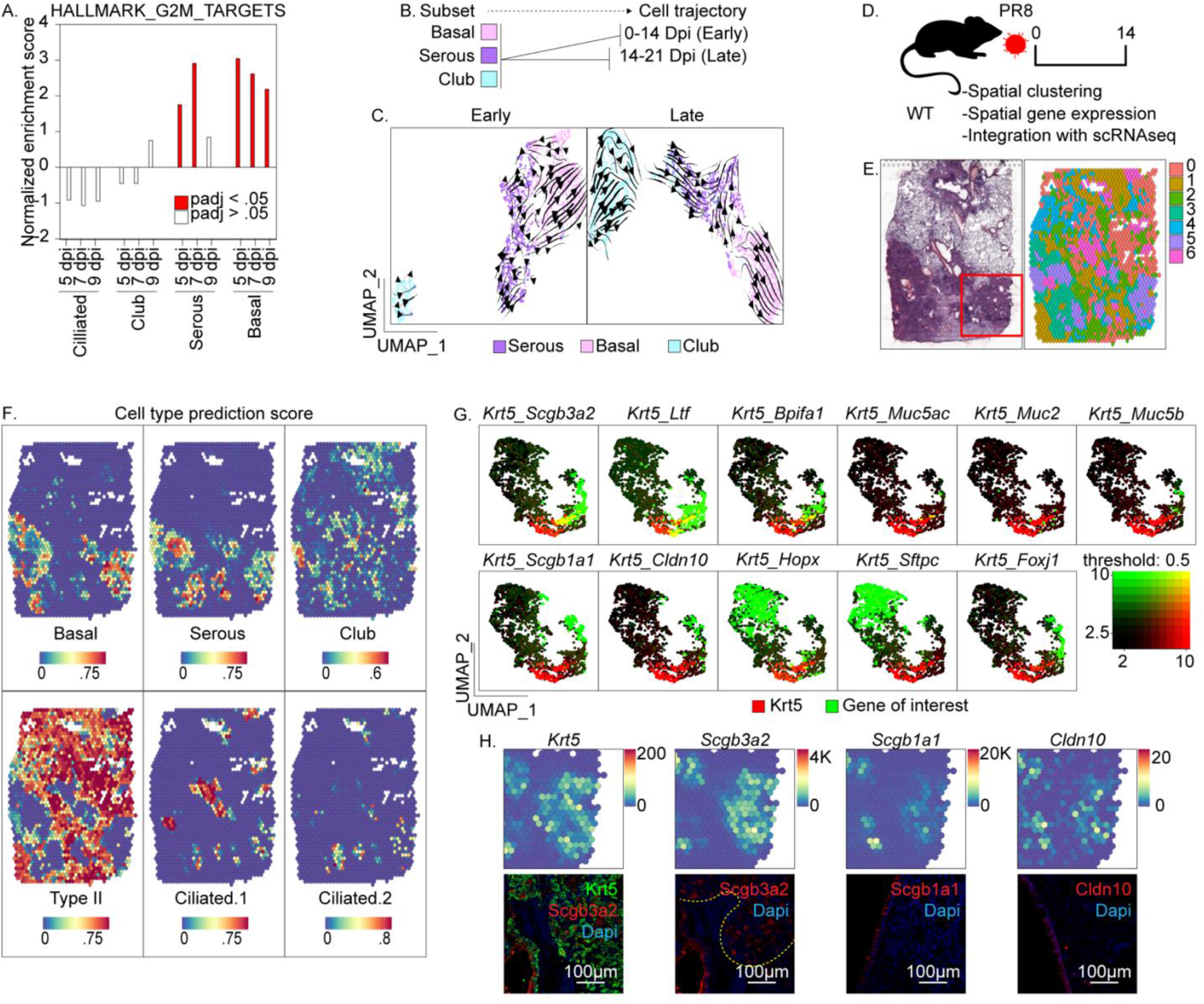
Single cell and spatial gene expression profiling to investigate lineage relationships between serous and basal cells following PR8 infection. (A) Comparative analysis of normalized enrichment for mitotic gene sets by cell type & condition. Red bars represent conditions with adjusted p value <0.05. (B) Experimental design for generation of RNA velocity projections. Single cell RNAseq data were first subsetted by cell type (basal, serous, and club), followed by time following infection (early: naïve, 3, 5, 7, 9, 11, 14 dpi, or late: 14, 17, and 21 dpi). (C) Trajectory inference based upon RNA velocity profiling among basal, serous and club cell subsets. (D) Experimental design for generation of spatial gene expression data. (E) Projection of spot clusters onto H&E image of tissue sample. (F) Cell type prediction scores of epithelial populations represented within spatial RNAseq data. Cell type specific gene signatures were generated using scRNAseq data in Fig. 1 A. Color scale at bottom reflects intensity of cell type prediction scores. (G) Co-expression of selected epithelial markers 14 days following PR8 exposure: *Krt5* (basal), *Scgb3a2* (serous and club), *Ltf, Bpifa1* (serous), *Muc5ac, Muc2, Muc2*(goblet), *Scgb1a1, Cldn10* (club), *Hopx* (Type I), *Sftpc* (Type II), *Foxj1* (Ciliated). (H) Spatial gene expression and corresponding immunofluorescence of cell type-specific markers. Transcripts were mapped onto spot coordinates from region sampled in red box in Fig. 2E: *Krt5* (basal), *Scgb3a2* (serous and club), *Scgb1a1* (club) and *Cldn10* (club) markers. Immunofluorescence colocalization of either Cldn10, Scgb1a1, or Scgb3a2 (red), with Krt5 (green), within injured distal airway of 14 dpi PR8 infected mice. Color scale reflects abundance of indicated transcripts.

We next used Visium spatial RNA-Seq to define the relationship between hBC and IS cells within lung tissue 14 days after PR8 infection (Fig. 2D). Unsupervised clustering revealed 7 transcriptionally distinct gene signatures that were projected onto a spatial map of the sampled tissue section (Fig. 2E). Cell type signature scores generated from scRNAseq (Fig. 1C) were applied to spatial RNAseq data to colocalize cell types (Fig. 2F). The resulting spatial maps of cell types demonstrated close spatial association of IS but not club cells with hBC’s, as suggested from the predicted lineage relationships between these cell types by RNA velocity analysis (Fig. 2F, G; Supp. Fig. 2B). We corroborated spatial RNAseq data by immunofluorescent localization of Scgb3a2 to regions of Krt5-immunoreactivity within remodeled regions of alveolar epithelium of PR8-infected mice (Fig. 2H).

### Scgb3a2^pos^Scgb1a1^neg^ IS cells demonstrate serous-basal-serous plasticity during repair following PR8

To validate our earlier predictions that IS but not club cells could serve as precursors for hBC in response to PR8 infection, we developed a lineage tracing approach to independently tag and fate map these secretory cell types. Our single-cell and spatial RNAseq data indicated that both IS cells and club cells express Scgb3a2 but that only club cells express Scgb1a1. Accordingly, we generated dual recombinase (DR) mice harboring a Dre/Cre reporter allele in conjunction with *Scgb1a1-CreER* and *Scgb3a2-DreER* recombinase driver alleles (Fig. 3A). Tamoxifen (TM) exposure of these mice results in Dre-mediated excision of a poly A signal (STOP) within Scgb3a2-expressing IS and club cells, with subsequent Cre-mediated excision of tdT-STOP within Scgb1a1-expressing club cells.

**Fig. 3.**
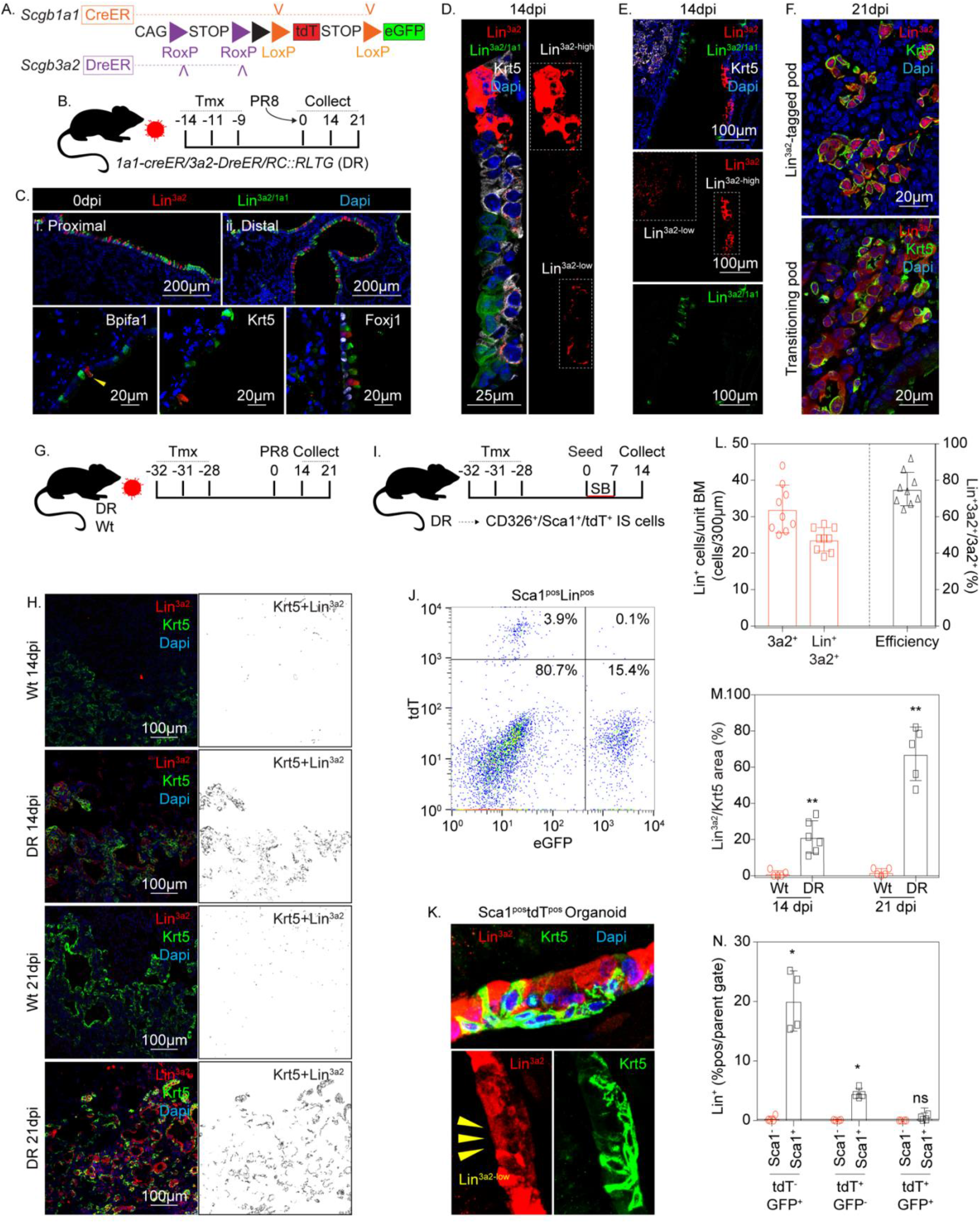
Rare Scgb3a2^+^/Scgb1a1^-^ serous cells reside within intralobar epithelium and assume BC fates in response to PR8-induced airway injury. (A) Schematic illustration of recombinase driver and reporter alleles used in Dre/Cre recombinase (DR) mice for independent lineage labeling of serous and club cells. (B) Experimental design. DR mice (n = 5 per experimental group) were treated with 3 doses of TM, infected with PR8 and recovered for the indicated time points. (C) Representative immunofluorescence localization of tdT (Lin^3a2^; red) and eGFP (Lin3^a2/1a1^; green) lineage reporters reflective of serous and club cells, respectively. Shown is a low magnification image (left) and selected high magnification images of either proximal (i) or distal (ii) airway epithelium (right). Bottom row shows representative immunofluorescence colocalization of lineage reporters (green or red) with either Bpifa1, Foxj1, or Krt5 (white) in airways of naïve mice. (D) Representative immunofluorescence colocalization of lineage reporters (green or red) with Krt5 (white) 14 days after PR8 infection, demonstrating Lin^3a2-high^ and Lin^3a2-low^ epithelial cells. (E) Representative immunofluorescence colocalization of lineage reporters (green or red) with Krt5 (white) 14 days after PR8 infection, demonstrating recruitment of Lin^3a2^ -low epithelial cells into damaged alveolar epithelium. (F) Representative immunofluorescence colocalization of tdT (Lin^3a2^; red) with Krt5 (green) among lineage-positive alveolar clusters 21 days after PR8 infection. (G) Experimental design for assessing lineage tagged populations using extended washout period. DR mice (n = 5 per experimental group) were treated with 3 doses of TM, infected with PR8 and recovered for the indicated time points. (H) Representative immunofluorescence colocalization of lineage reporters (red) with Krt5 (green) 14 and 21 days after PR8 infection using extended washout periods for TMX. (I) Experimental design for generation of in-vitro organoids from tdT-lineage tagged cells (n = 5). (J) Representative scatter plot of tdT and eGFP lineage labeled cells from tissue homogenate of DR mice. Lineage reporter (tdT; eGFP) was assessed as a function of surface expression of CD326^+^Sca1^+^ airway epithelium and CD326^+^Sca1^-^ alveolar epithelium. (K) Representative immunofluorescence colocalization of lineage reporters (Lin^3a2^; red) with Krt5 (green) from organoid cultures demonstrating Lin^3a2-high^ and Lin^3a2-low^ epithelial cells. (L) Quantification of recombination efficiency following immunofluorescent staining for tdT, eGFP and Scgb3a2 (as in ‘D’; TM washout period = 9 days). Red bars show either Scgb3a2 or lineage traced (either tdT or eGFP) cells per unit basement membrane. Black bar represents the fractional representation of Scgb3a2-immunofluoresecnt cells that are lineage positive. (M) Contribution of lineage-tagged populations to hBC in alveolar ‘pods’ as a function of time (as in ‘H’ ; TM washout period = 28 days) after PR8 infection (G). N=5 per condition, with significance determined by Mann-Whitney U-test. (N) Quantification of flow cytometry (as in ‘J’ ; TM washout period = 28 days) from tissue homogenate from PR8-infected *DR* mice. Lineage reporter (tdT; Lin^+^) was assessed as a function of surface expression of Sca1 and data presented as the fraction of positive cells.

Outcomes of these recombination events are tracing of Scgb3a2^+^/Scgb1a1^-^ IS cells by expression of tdT (Lin^3a2^), and Scgb3a2^+^/Scgb1a1^+^ club cells by expression of eGFP (Lin^3a2/1a1^; Fig. 3A). For lineage tracing after PR8 infection, no differences were observed in the fate of either IS or club cells with 9 day or 28 day TM washout periods. Consequently, 9 or 28 day TM washout periods were used interchangeably for PR8 recovery experiments. Tamoxifen exposure was followed by a wash-out period of 9 days before analysis of lineage tracing in lungs of naïve mice (Fig. 3B). Control experiments involving TAM exposure of mice harboring *Scgb3a2-DreER* and Dre/Cre reporter but lacking *Scgb1a1-CreER*, revealed efficient induction of tdT within Scgb3a2-expressing airway cells without evidence of eGFP reporter expression, confirming the specificity of DreER for RoxP without recombination of LoxP sequences (Supp. Fig. 3A-C). We went on to assess the steady-state phenotype of both IS and club cells in dual recombinase mice harboring *Scgb3a2-DreER, Scgb1a1-CreER*, and Dre/Cre recombinase substrate alleles. Lineage labeling of Scgb3a2 immunoreactive cells in airways of naïve mice was determined to be 75.2 ± 3.06%, of which Lin^3a2^ IS cells (16.8% of total lineage-labeled cells) were interspersed among Lin^3a2/1a1^ club cells (83.2% of total lineage-labeled cells) (Fig. 3C, D, L). Lineage-labeled (tdT^+^) serous cells in airways of naïve mice were the only cell type that showed positive immunofluorescent staining for Bpifa1, and were uniformly negative for markers of ciliated and BC (Fig. 3D.). Notably, evidence linking IS cells to other proposed progenitors of hBC was tenuous (Supp. Fig. 3D-G).

We next sought to assess the fate of lineage-labeled IS and club cells following PR8-induced lung injury. At the 14 day post-PR8 recovery time point, both Lin^3a2/1a1^ and Lin^3a2^ cells were observed in airways (Fig. D, E). Whereas Lin^3a2/1a1^ cells occupied a luminal location and were exclusively Krt5^-^, Lin^3a2^ cells occupied both luminal and basal locations with basal-localized cells showing positive Krt5 immunoreactivity. The observed behavior of Lin^3a2/1a1^ cells is consistent with prior reports by others and us that club cells do not contribute significantly to the expanding pool of BC in response to lung injury or infection (Ray et al., 2016; Vaughan et al., 2015). Interestingly, Lin^3a2^ cells included both tdT-bright (Lin^3a2-high^) and tdT-dim (Lin^3a2-low^) populations, this property correlating with Krt5^-^ and Krt5^+^ immunoreactivity, respectively. Injured alveolar regions were repopulated exclusively by basal-like Krt5-immunoreactive cells that were Lin^3a2-low^ at day 14 post-infection (Fig. 3E). By recovery day 21, Lin^3a2^ cells occupying alveolar regions included Krt5^+^Lin^3a2-low^, Krt5^+^Lin^3a2-high^, and Krt5^-^ Lin^3a2-high^ patches, with some patches appearing to show mixed phenotypes suggestive of transitioning cell states (Supp. Fig. 3H, I; Fig. 3F).

Control experiments with an extended washout period (28 days) were performed to ensure no residual tamoxifen-mediated recombination that could confound interpretation of lineage tracing experiments. Results with a 28 day washout mirrored those using a 9 day washout, suggesting that washout periods of 9 days or greater, under the conditions of tamoxifen preparation and delivery used in this study, were sufficient to faithfully assess IS/club cell fate after PR8 infection (Fig. 3G, H, M). Immunofluorescence staining for Scgb3a2 confirmed that cells phenotypically similar to IS cells were included among the alveolar Krt5^-^Lin^3a2-high^ population (Supp. Fig. 3J). Furthermore, hBC did not show clear evidence of a differentiation trajectory towards alveolar epithelium (Supp. Fig. 3K, L). These data are consistent with trajectory analysis of scRNAseq data suggesting that hBC observed after PR8 infection eventually yield differentiating airway epithelial cells such as Scgb3a2-expressing IS cells (Fig. 2C). Expanding Lin^3a2-high^ cells displayed characteristics of IS cells and show increasing abundance with time of recovery after PR8 infection that validate observations made by scRNAseq (Fig. 1E).

To further verify that both Lin^3a2-high^ and Lin^3a2-low^ epithelial cells were derived from Lin^3a2-high^ IS cells, we FACS enriched Lin^3a2^ IS cells and evaluated their clonal potential in 3D organoid assays (Fig. 3I, J, N). Consistent with the observed behavior of Lin^3a2-high^ IS cells in vivo, IS-derived organoids recapitulated the pseudostratified epithelial structure of conducting airways including both Lin^3a2-high^/Krt5^-^ and Lin^3a2-low^/Krt5^+^ epithelial cells (Fig. 3K). Our data confirm that both Lin^3a2-high^ and Lin^3a2-low^ cells are derived from a common Lin^3a2-high^ IS progenitor, and that changes in reporter fluorescence intensity correlate with alterations in cell state. Taken together, the changing milieu of the PR8 injured lung initially results in specification of hBC from an airway IS progenitor, followed by their differentiation back to IS and other luminal cell types in airways and injured alveolar regions.

### Regulatory gene networks involved in immune-epithelial crosstalk are activated in nascent basal but not in IS cells

Next, we sought to define mechanisms that regulate specification and fate of hBC observed in lungs of PR8 infected mice. To gain further insights into pathways regulating the behavior of either IS or BC, we evaluated changes in regulatory genes over the time course of PR8 infection and recovery using Bigscale2 (Iacono et al., 2019; Iacono et al., 2018). ScRNAseq data from naïve and days 11-17 after PR8 infection were sorted by cell type to yield four regulatory gene networks (GRN; Supp. Fig. 4A). Cell type specific GRNs from post-infection time points were then compared to their respective naïve cell controls to normalize node and edge values (Supp. Fig. 4B). A gene list of the top 300 delta degree node centralities was generated and used to identify enriched gene ontology (GO) terms by Panther-based overrepresentation tests (Supp. Fig. 4C; Fig. 4A). Gene regulatory networks shown in Supp. Fig. 4C demonstrate dynamic changes occurring among BC but less so for IS cells between naïve and post-infection time points. Highly enriched GO terms in basal populations included pathways involving cell proliferation, such as activation of canonical Wnt signaling (Haas et al., 2019) (not shown), consistent with observed BC proliferation (Fig. 2A) and associated hyperplasia (Fig. 1D) that accompanies recovery from PR8 infection. Pathways associated with cytokine signaling and innate immune stimulation of epithelial cells were also significantly up regulated in BC, and to a lesser extent in IS cells, isolated from PR8 infected lungs (Fig. 4A).

**Fig. 4.**
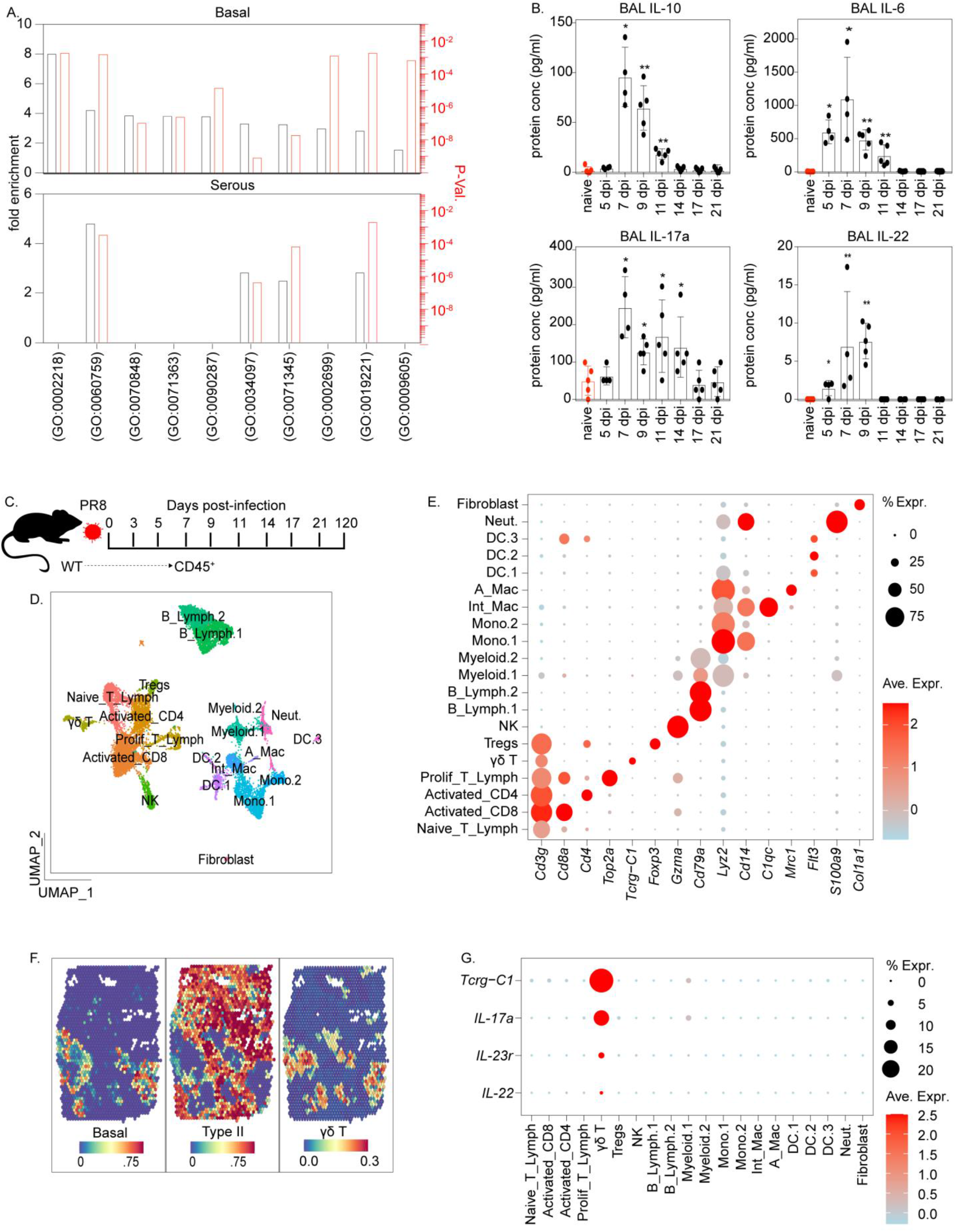
T lymphoid cells colocalize with hBC at sites of lung injury following PR8 infection. (A) Representation of significant GO terms associated with immune>epithelial interaction enriched among PR8-infected BC (upper) and IS cells (lower) using a Panther overrepresentation test. (B) Assessment of selected cytokine levels in BALF during recovery from PR8 infection. 4-5 biological replicates were used per timepoint with statistical significance determined by Mann-Whitney U-test. (C) Experimental design. Mouse lung homogenates were collected from 8-12-week-old C57/Bl6 mice. CD31^-^CD326^+^CD45^+^ immune cells were enriched by FACS and transcriptomes assessed by scRNAseq (n = 5 per group). (D) UMAP plot of combined scRNAseq data. Unsupervised clustering was used to distinguish distinct cell phenotypes which were assigned to known immune cell types based upon gene signatures. (E) Dot Plot selected cell-type specific gene expression for each annotated cluster. (F) Cell type prediction scores of immune populations represented within spatial RNAseq data. Cell type-specific gene signatures were generated using immune scRNAseq data in Fig. 5A and epithelial scRNAseq data in Fig. 1A. Color scale at bottom reflects intensity of cell type prediction scores. (G) Dot plot showing activation of type-17 gene signature(IL-17a, IL-23r and IL-22) in γδT cells

Innate immune activation is a well-documented response to respiratory viral infection in general and is a key regulator of epithelial cell fate leading to remodeling in airways and increased postnatal susceptibility to allergic inflammation (Hackett et al., 2011). Notably, local production of TNFα and IL-1β following influenza virus infection in mice promote the regenerative capacity of alveolar epithelium (Katsura et al., 2019), and IL-22 enhances survival and reduces lung fibrosis following PR8 infection (Pociask et al., 2013). To explore further the potential roles for cytokine-mediated changes in epithelial cell fate, we performed a multiplex protein assay to quantify cytokine responses in lungs of influenza virus infected C57/Bl6 mice. We observed a significant increase in interferon (Ifnγ) and pro-inflammatory cytokines IL-6, TNFα and IL-1β (Fig. 4B; Supp. Fig. 4D), consistent with known responses to respiratory viral infection (Katsura et al., 2019). A significant induction of type 17-related cytokines, including IL-17a and IL-22, was observed in lung tissue homogenates and/or bronchoalveolar lavage (BAL) of mice between recovery days 5-11 after PR8 infection (Fig. 4B, Supp. Fig. 4D). The protective effects of IL-22 (Pociask et al., 2013) and the coincidence of increased cytokine production with BC expansion in airways and alveoli led us to speculate that IL-22 signaling plays a role in BC expansion following PR8 infection.

### Renewal and differentiation of hBC is regulated by γδT-cell derived IL-22

To further explore roles for innate immune activation and IL-22 in regulation of epithelial cells in the airways and alveoli of PR8 exposed mice, we used scRNAseq to define changes to immune cell populations elicited in response to infection (Fig. 4C). ScRNAseq data were collected from CD45^+^ lung cells recovered at different times following PR8 infection. UMAP dimensional reduction of aggregated data allowed visualization of major immune subsets (Fig. 4D, E). We sought to define spatial context of these major immune subsets in relation to hBC observed during recovery. Gene signatures derived from immune enriched scRNAseq dataset (Fig. 4D) were used to infer cell localization within spatial RNAseq data (Fig. 4F). Spatial gene expression analysis indicates preferential colocalization of γδT cells within BC-rich regions in recovering lung tissue (Fig. 4F). Interestingly, type 17 γδT cells show increased abundance within regions of BC hyperplasia (Fig.4F, G). This led us to consider the possible regulatory influence of γδT lymphoid subset over BC fate. Since several immune subsets can secrete IL-22, we generated mice harboring *IL-22*^*Cre*^/*ROSA-26-tdT* to fate map IL-22-lineage immune cells during the repair response to PR8 infection (Fig. 5A). To identify the cell types expressing IL-22, we gated IL-22 lineage tag (tdT) CD45^+^ cells for expression of lymphoid markers CD3 and CD4 (Fig. 5B). We found that CD3^+^CD4^-^ T cells subsets represented the bulk of IL-22-lineage immune cells elicited in response to PR8 infection, with minor contributions made by CD3^+^CD4^+^ Th17 cells (Fig. 5B, Supp. Fig. 5A). Non-T lymphoid CD3^-^ subsets did not contribute to the IL-22-expressing lineage following PR8 infection (Supp Fig. 5A). Immunofluorescence analysis demonstrated that IL-22 and γδTcr-immunoreactive immune cells localize to hBC and were induced in response to acute lung injury (Fig. 5C-E).

**Fig. 5.**
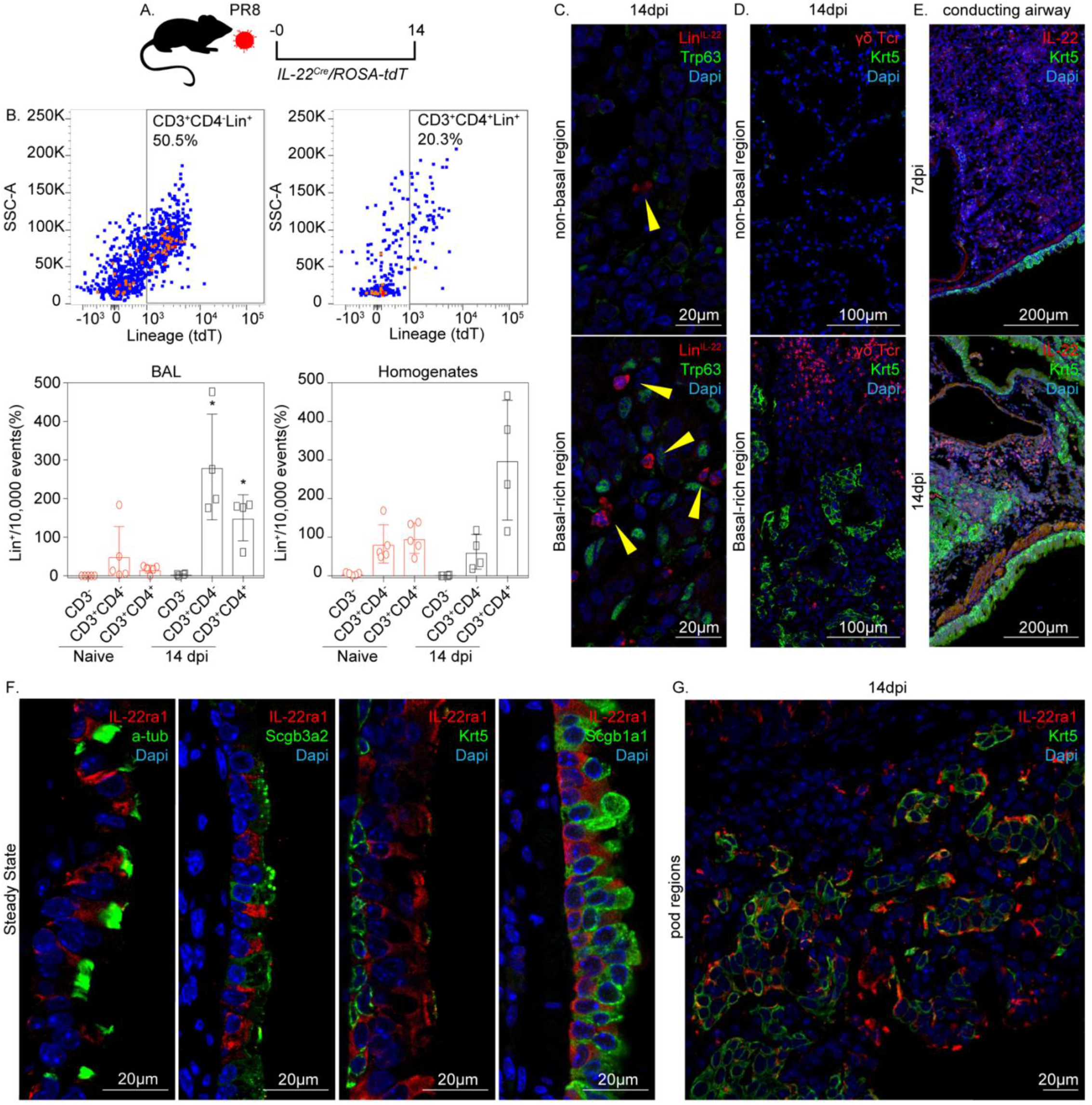
Identity of IL-22 expressing cells and their localization to regions of BC hyperplasia in lungs of PR8 infected mice. (A) Experimental design for fate mapping of IL-22 expressing cells following PR8 infection using *IL-22*^*Cre*^/*ROSA-tdT* mice. (B) Flow cytometry analysis of either BAL or tissue homogenate from PR8-infected *IL-22*^*Cre*^/*ROSA-tdT* mice. IL-22 lineage reporter (tdT; Lin^+^) was assessed as a function of surface expression of CD3 and CD4 and data presented as the fraction of positive cells. (C) Immunofluorescence detection of Lin^+^ cells as a function of p63-immunoreactive BC within BC-rich vs BC devoid alveolar epithelium of PR8-infected mice. (D) Immunofluorescence detection of γδTcr^+^ cells as a function of Krt5-immunoreactive BC within BC-rich vs BC devoid alveolar epithelium of PR8-infected mice. (E) Immunofluorescence localization of IL-22 expressing cells 7 and 14 days post-PR8 infection. (F) Immunofluorescence colocalization of IL-22ra1 (red) with either a-tub (ciliated), Scgb3a2 (serous/club), Krt5 (BC), Scgb1a1 (club) (green) in airways of naïve mice. (G) Representative immunofluorescent colocalization of IL-22ra1 (red) and Krt5 (green) in BC-rich alveolar region of PR8-infected mouse lung.

To identify epithelial cell types capable of responding to IL-22 signaling, we use immunofluorescence to identify which epithelial subsets express IL-22ra1, the subunit of heterodimeric IL-22 receptor that confers ligand specificity to IL-22. Notably, within the conducting airway of naïve mice, the highest level of IL-22ra1 immunoreactivity was observed among post-mitotic ciliated epithelial cells, with little to no immunoreactivity seen on club, basal or serous cells (Fig. 5F). However, a significant level of IL-22ra1 expression was observed in alveolar hBC (hereon called pods (Kumar et al., 2011)) during recovery following PR8 infection (Supp. Fig. 6A; Fig. 5G). Interestingly, the intensity of IL-22ra1 staining was greatest within the outer periphery of pod regions indicating altered basal responsiveness to IL-22 during injury (Supp. Fig. 6B). Taken together, our data suggest a process of altered IL-22 responsiveness exclusively among nascent basal populations of the PR8-injured lung that regulate BC fate.

### IL-22 promotes self-renewal of hBC, allowing hyperplastic expansion in airways and injured alveoli

To investigate the contribution made by IL-22 in regulation of BC fate after PR8 infection, we used *IL-22*^*Cre/Cre*^ (IL-22 LOF) and *IL-22ra1*^*fl/fl*^/*Shh-Cre* (IL-22r cLOF) mice to modulate IL-22 levels and signaling, respectively (Fig. 6A). Compared to their corresponding WT control groups, neither IL-22 LOF nor IL-22r cLOF mice showed significant changes in weight loss, viral gene expression, or loss of parenchymal Pdpn immunofluorescence, following PR8 infection (Supp. Fig. 7A, B, C; Fig. 6B). Next, we assessed expansion of Krt5-immunoreactive BC as a function of damaged area in IL-22 LOF mice following PR8 infection. A reduction in Krt5^+^ BC hyperplasia was observed in IL-22 LOF mice compared to WT control mice at all time points examined, which reached statistical significance by the day 17 recovery time point (Fig. 6B, D). These data were confirmed by assessing *Krt5* mRNA content within total lung RNA isolated at each timepoint, for which statistically significant declines in *Krt5* mRNA were observed at both 14 day and 17 day recovery time points (Fig. 6E). Similar observations of reduced BC expansion were seen among IL-22r cLOF mice following PR8 infection (Supp. Fig. 7C).

**Fig. 6.**
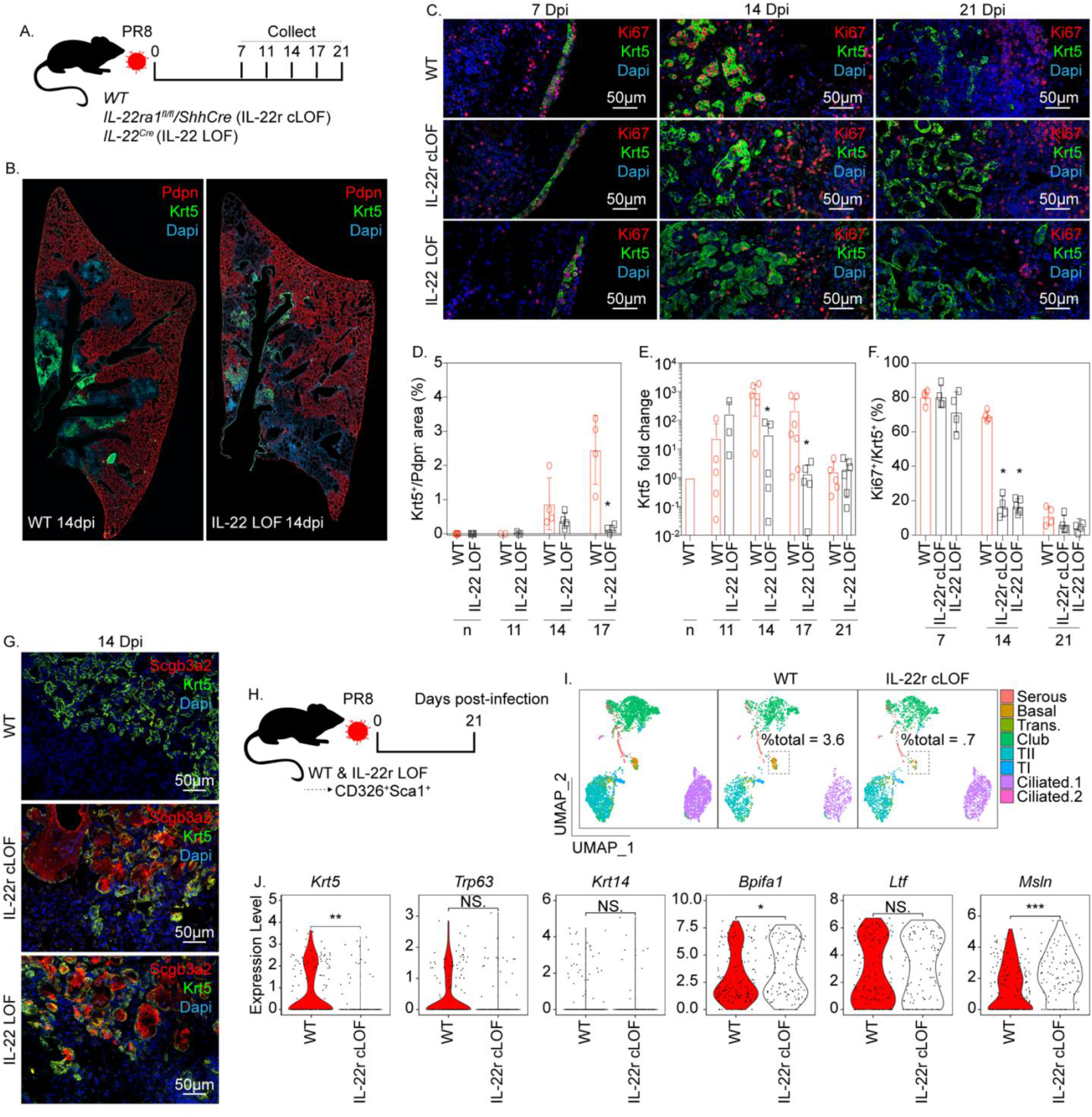
IL-22 regulates BC fate in the PR8-injured lung. (A) Experimental design to assess BC expansion in *IL-22*^*Cre*^ homozygous (IL-22 LOF) mice. Left lobes were collected for immunostaining and total RNA isolated from the right cranial lobe for gene expression analysis. (B) Representative immunofluorescence localization of Pdpn (red) and Krt5 (green) in lungs of WT and IL-22 LOF mice 14 days post-PR8 infection. (C) Representative immunofluorescence localization of Ki67 (red) and Krt5 (green) in lung tissue of PR8-infected WT, IL-22 LOF and *IL-22ra1*^*fl/fl*^/*Shh-Cre* (IL-22r cLOF) mice. (D) Quantification of (B). The fraction of Krt5 stained area was normalized to the area of damaged alveolar epithelium (DAPI^+^Pdpn^-^) (n = 3-5 biological replicates per group). Statistical analysis was performed by Mann-Whitney U-test. (E) qPCR detection of *Krt5* mRNA in total lung RNA of PR8-infected WT and IL-22 LOF mice. 3-5 biological replicates were used per condition. Statistical analysis was performed by Mann-Whitney U-test. (F) Quantification of (C). Shown is the Ki67-labeling index for BC, expressed as percentage of Ki67^+^, Krt5^+^ cells within areas of BC hyperplasia (n = 3-5 biological replicates per group). Statistical analysis was performed by Mann-Whitney U-test. (G) Representative immunofluorescence localization of Krt5 (green) and Scgb3a2 (red) in lungs of PR8-infected WT, IL-22 LOF and IL-22r cLOF mice. (H) Experimental design for scRNAseq of PR8-infected IL-22r cLOF mice (n = 5 biological replicates per group). (I) UMAP plot of Sca1^+^ airway enriched cells from IL-22r cLOF and C57/Bl6 control. Cell type annotation of clusters was performed based upon expression of cell type-specific genes. UMAP plot were re-clustered based on genetic permutation to better represent changes in BC populations between experimental conditions. (K) Assessment of selected basal (*Krt5, Krt14, Trp63*) and serous (*Bpifa1, Ltf*,, *Msln*) genes between IL-22r cLOF and WT control.

Similarly, in other tissues and carcinomas, IL-22 regulates the proliferative potential of epithelial stem progenitors and tumor cells (Jin et al., 2019). To assess if reduced BC hyperplasia observed in IL-22 LOF mice was associated with reduced epithelial proliferation, we measured the BC proliferative index by immunofluorescence of Ki67 and Krt5 during recovery following PR8 infection. The Ki67-labeling index of BC observed at 7 days post PR8 infection did not differ between IL-22 LOF, IL-22r cLOF, and their corresponding WT controls (Fig. 6C, F). These data indicate that BC expansion at early time points after PR8 infection occurs in an IL-22-independent manner and are consistent with the observed lack of IL-22ra1-immunoreactivity in airway BC of either naïve control mice or mice at early time points after PR8 infection (Fig. 5F). However, hBC observed within airway and alveolar regions of both IL-22 LOF and IL-22r cLOF mice at 14 days post-PR8 infection showed significantly reduced levels of Ki67 staining compared to their corresponding WT controls (Fig. 6C, F). These data indicate that at later recovery time points, IL-22 promotes BC expansion and self-renewal, resulting in maintenance of proliferative potential. All genotypes showed reduced BC Ki67 proliferative indices that did not differ significantly between groups at 21 days post-PR8 infection (Fig. 6C, F). To determine if the lack of IL-22 stimulation impacts the fate of alveolar BC during recovery from PR8 infection, we co-stained pod regions observed in parenchymal tissue of PR8-infected WT and IL-22 LOF mice for BC and IS markers. Interestingly, Krt5-immunoreactive pods observed in lungs of either IL-22 LOF or IL-22r cLOF mice 14 days post-PR8 infection displayed evidence for Scgb3a2 expression that was absent in comparable Krt5-immunoreactive pod structures of wildtype control mice (Fig. 6G). The appearance of pod structures composed of epithelial cells showing colocalization of both Krt5 and Scgb3a2 immunofluorescence is consistent with earlier data showing BC>IS differentiation, and supports the notion that residual Krt5-immunoreactivity reflects the BC origin of differentiating IS cells. To further examine the impact that loss of IL-22 has on BC maturation, scRNAseq profiles were generated for Sca1^+^ airway enriched cells isolated from lungs 21 days after PR8-infection of either IL-22r cLOF mice or their corresponding WT controls (Fig. 6H). Reduced BC numbers in lungs of IL-22r1a-cLOF mice compared to WT control mice was confirmed by UMAP clustering (Fig. 6I). A significant decrease in the abundance of epithelial cells expressing BC marker genes, was observed among IL-22r1a-cLOF mice compared to their WT controls (Fig. 6J). Collectively, our data suggest that IL-22 functions to promote BC renewal over differentiation and that down regulation of innate immune responses at late time points following PR8 infection with associated reduction in IL-22, serves as a trigger to promote basal to serous cell differentiation.

## Discussion

Basal cell (BC) hyperplasia and colonization of the injured alveolar gas-exchange region are significant determinants of tissue remodeling and morbidity among patients with severe respiratory viral infections and share features of distal lung remodeling in patients with interstitial lung disease. Here we show that hBC elicited by infection of the mouse respiratory tract with H1N1 influenza virus (strain PR8) are derived predominantly from a serous cell subset of intralobar secretory cells. IiBC were distinguished from pre-existing BC by their relatively immature molecular phenotype and expression of IL-22Ra1. Innate immune activation and associated secretion of IL-22 promoted self-renewal of hBC in airways and establishment of hyperplastic foci within injured alveoli. Germline loss of IL-22 or conditional loss of IL-22ra1 expression within epithelial cells, limited expansion and promoted premature differentiation of hBC into serous cells. These findings establish a novel mechanism that promotes expansion of serous cells and their colonization of injured alveolar epithelium, leading to epithelial remodeling and loss of normal alveolar epithelium following respiratory viral infection.

We provide evidence for the existence of two independent secretory cell lineages within intralobar airways of the mouse lung, club cells and intralobar serous (IS) cells. Previous lineage tracing studies have demonstrated that Scgb1a1-expressing club cells are capable of unlimited self-renewal and replacement of specialized epithelial cell types of bronchiolar airways in mice (Rawlins et al., 2009). However, epithelial cell injury in lungs of PR8 infected mice leads to activation of non-canonical epithelial progenitors leading to BC hyperplasia (Ray et al., 2016; Vaughan et al., 2015; Xi et al., 2017; Yang et al., 2018; Zuo et al., 2015). We show that only the IS subset of secretory cells can assume BC fates in the setting of severe respiratory viral infection, suggesting that IS cells function as a reserve epithelial progenitor that are unlikely to make significant if any contribution to homeostatic epithelial maintenance in distal airways. Expansion of IS cells following influenza virus infection and their phenotypic conversion to hBC, suggest an unappreciated role for IS cells in epithelial maintenance and repair following severe injury. Interestingly, IS cells labeled using the dual recombinase lineage-tracing approach outlined in our study showed no evidence of p63 immunoreactivity, suggesting that their contribution to basal cell expansion following PR8 infection are distinct from those of rare p63-lineage epithelial cells described in lineage tracing studies using a *Trp63-CreER* driver allele (Xi et al., 2017; Yang et al., 2018). In our unpublished studies we find evidence of reduced basal cell expansion in *Trp63-CreER* mice, suggesting that haploinsufficiency of p63 isoforms impacts either specification or expansion of nascent basal cells following PR8 infection. Interestingly, IS cells were among a heterogeneous population of H2-k1 high epithelial cells observed in lungs of either naïve or PR8-infected mouse lungs, raising the possibility that they represent a more defined subset of H2-k1 cells proposed as BC progenitors (Kathiriya et al., 2020).

Our findings also shed new light on the biological significance of earlier work by Tata et al., who demonstrated that SSEA^+^ secretory cells of tracheobronchial airways have the unexpected capacity to replenish basal stem cells in a genetic model of BC ablation (Tata et al., 2013). However, even though IS cells identified in our study function in a similar capacity to SSEA^+^ secretory cells defined by Tata et al., to yield hBC, we show that the fate/stemness of IS-derived BC is dictated by the regional microenvironment. Expansion of IS-derived hBC and subsequent differentiation led to proximalization of distal conducting airways through replacement of the normally simple cuboidal bronchiolar epithelium with a pseudostratified epithelium, thus demonstrating their multipotency in the airway microenvironment like that described for SSEA^+^ secretory cell-derived BC in the Tata et al., study. Furthermore, hBC colonizing injured alveolar regions underwent significant proliferative expansion leading to localized BC hyperplasia, followed by differentiation into IS cells that replaced normal alveolar epithelium with a dysplastic serous cell predominant epithelial lining. It is possible that the observed influence of microenvironment on fate of IS-derived BC following PR8 infection may simply reflect the broader injury elicited by viral infection in our study compared to that resulting from targeted ablation of resident BC in tracheobronchial airways as in the study by Tata and colleagues. However, PR8 infection of mice is more akin to the type of lung injury seen in patients with respiratory viral infection and wide-spread lung injury that accompanies chronic lung disease. Importantly, our data provide insights into mechanisms of epithelial remodeling observed in distal lung tissue of patients with idiopathic pulmonary fibrosis, where alveolar epithelial progenitor cell dysfunction is associated with BC hyperplasia in small airways and establishment of dysplastic cysts in place of normal alveolar epithelium (Carraro et al., 2020; Seibold et al., 2013; Xu et al., 2016).

Even though hBC elicited by PR8 influenza virus infection share many properties of pre-existing BC of pseudostratified airways, their immature molecular phenotype and unique expression of IL-22ra1 impart distinctive functional properties that promote rapid epithelial replacement in airways and alveoli. We found that IL-22-expressing γδT-cells are recruited to sites of PR8-induced airway and alveolar injury, and that their local production of IL-22 promoted self-renewal of hBC with no impact on either their specification in airways or migration to injured alveoli. Notably, either germline loss of IL-22 or conditional loss of IL-22ra1 within lung endoderm resulted in premature differentiation of alveolar BC into Scgb3a2^+^ serous cells, thus limiting BC hyperplasia. These data shed new light on mechanisms of fibrosis in lungs of IL-22^-/-^ mice following PR8 infection (Pociask et al., 2013), where IL-22 restrains hBC in a highly proliferative and migratory state allowing expansion and re-epithelialization of injured airways and alveoli. We propose a model in which IS cells ultimately colonize injured alveoli following PR8 infection through a combination of IS>BC>IS phenotypic plasticity. Interestingly, these roles for IL-22 in regulating PR8-elicited BC in the lung are in contrast to observations made in other organ systems, such as in the epidermis where IL-22 regulates fibroproliferative responses associated with wound closure (McGee et al., 2013) and in the intestine, where IL-22 regulates epithelial cell fate and host defense through induction of antimicrobial factors (Lo et al., 2019; Zheng et al., 2008). We attribute this differences in outcome to target cells that respond to local production of IL-22 and to the impact of non-cell-autonomous influences of IL-22 signaling within epithelial and stromal cell types. Interestingly, in prior studies IL-22 has been shown to protect against influenza virus-induced pneumonia (Hebert et al., 2020) and has been implicated in the production of antimicrobial factors that protect against secondary bacterial subsequent to influenza virus infection (Abood et al., 2019). Our data demonstrate that these effects are an indirect consequence of IL-22-mediated expansion of IS-derived BC followed by their re-differentiation to yield IS hyperplasia and provide novel insights into mechanisms of protection against secondary bacterial pneumonia after respiratory viral infection.

## Acknowledgments

We would like to thank Matt Kostelny, Guangzhu Zhang and Katherine Drake for assistance with animal husbandry, and Stephen Beil for general laboratory support. We acknowledge support from the Applied Genomics, Pavilions Flow Cytometry & Cell Sorting, and Mouse Genetics cores at Cedars Sinai Medical Center (CSMC). This research was supported by grants from the National Institutes of Health (NIH) (R01 HL135163; P01 HL108793) to BRS, by the Bram and Elain Goldsmith Chair in Gene Therapeutics Research, and by the Office of Graduate Education at CSMC.

## Author contributions

Conceptualization, A.K.B., B.R.S.; methodology, A.K.B., J.Z., C.Y., G.C., E.I., A.L.C., B.R.S; data analysis, A.K.B., B.R.S; investigation, A.K.B., J.Z., C.Y., B.R.S; resources, E.I., A.L.C., C.M.H., J.K., B.R.S.; writing – original draft, A.K.B., B.R.S. ; writing –review & editing, A.K.B., J.Z., C.Y., G.C., E.I., A.L.C., C.M.H., J.K.K., W.C.P., B.R.S.; visualization, A.K.B., B.R.S.; supervision, A.K.B., J.Z., C.Y., G.C., W.C.P., B.R.S.; funding acquisition, B.R.S

---

**Supplemental Fig. 1.**
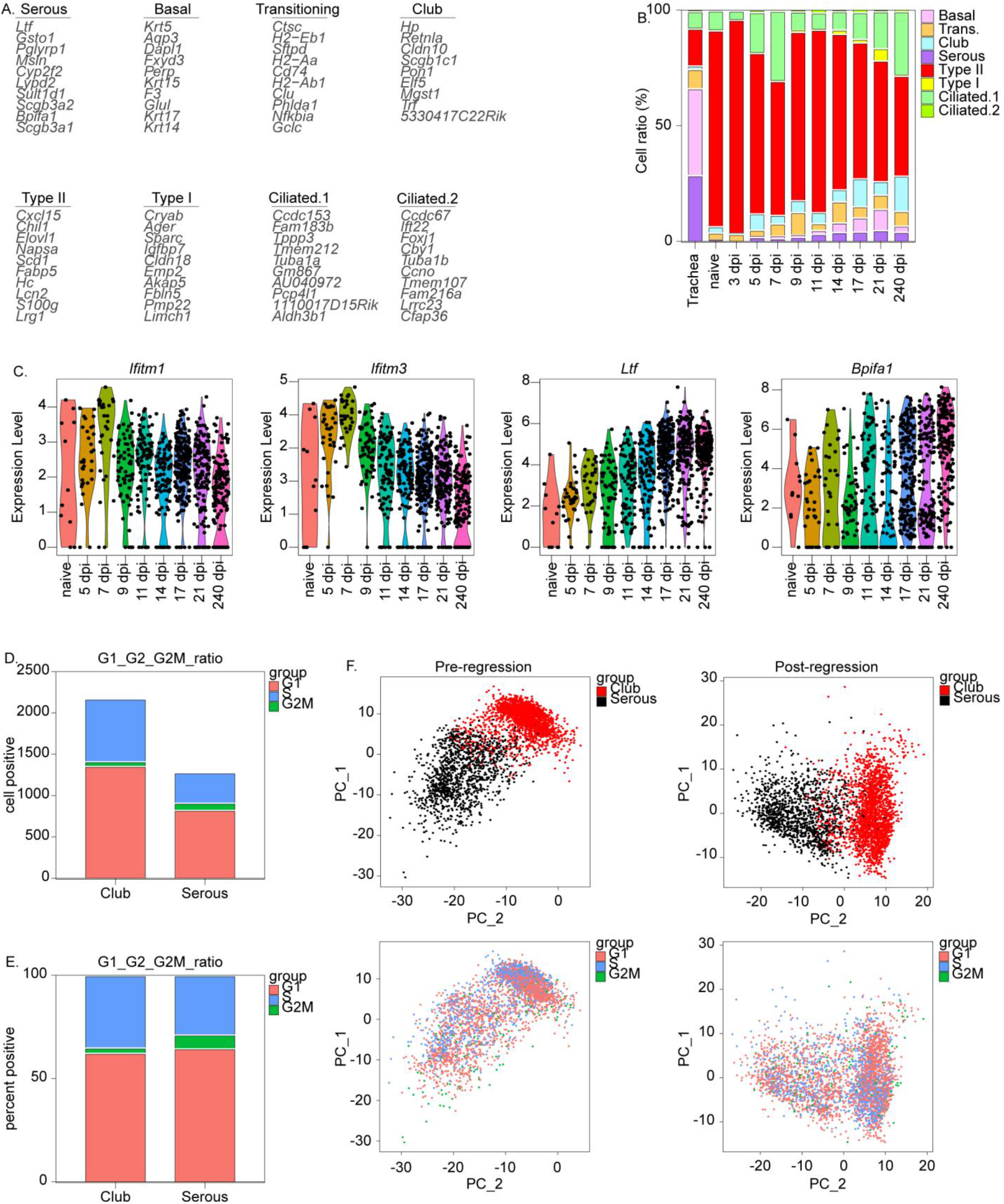
Club and IS cell cluster independently of cell cycle genes. (A) Full gene list for heatmap shown in Fig 1. A. (B). Representation of lung epithelial cell types as a function of percent total sampled cells at each recovery time point. dpi = days post infection. (C) Violin plot comparing expression of selected antimicrobial genes at indicated time points after PR8 infection. (D) Total G1_G2_G2M ratio between subsetted club and serous cells. (E) Percent G1_G2_G2M ratio between subsetted club and serous cells. (F) Principal component analysis between club and serous cells pre- & post-regression of cell cycle genes. Upper plots show points labeled by cell annotation and lower plots show points labeled according to G1_G2_G2M score.

**Supplemental Fig. 2.**
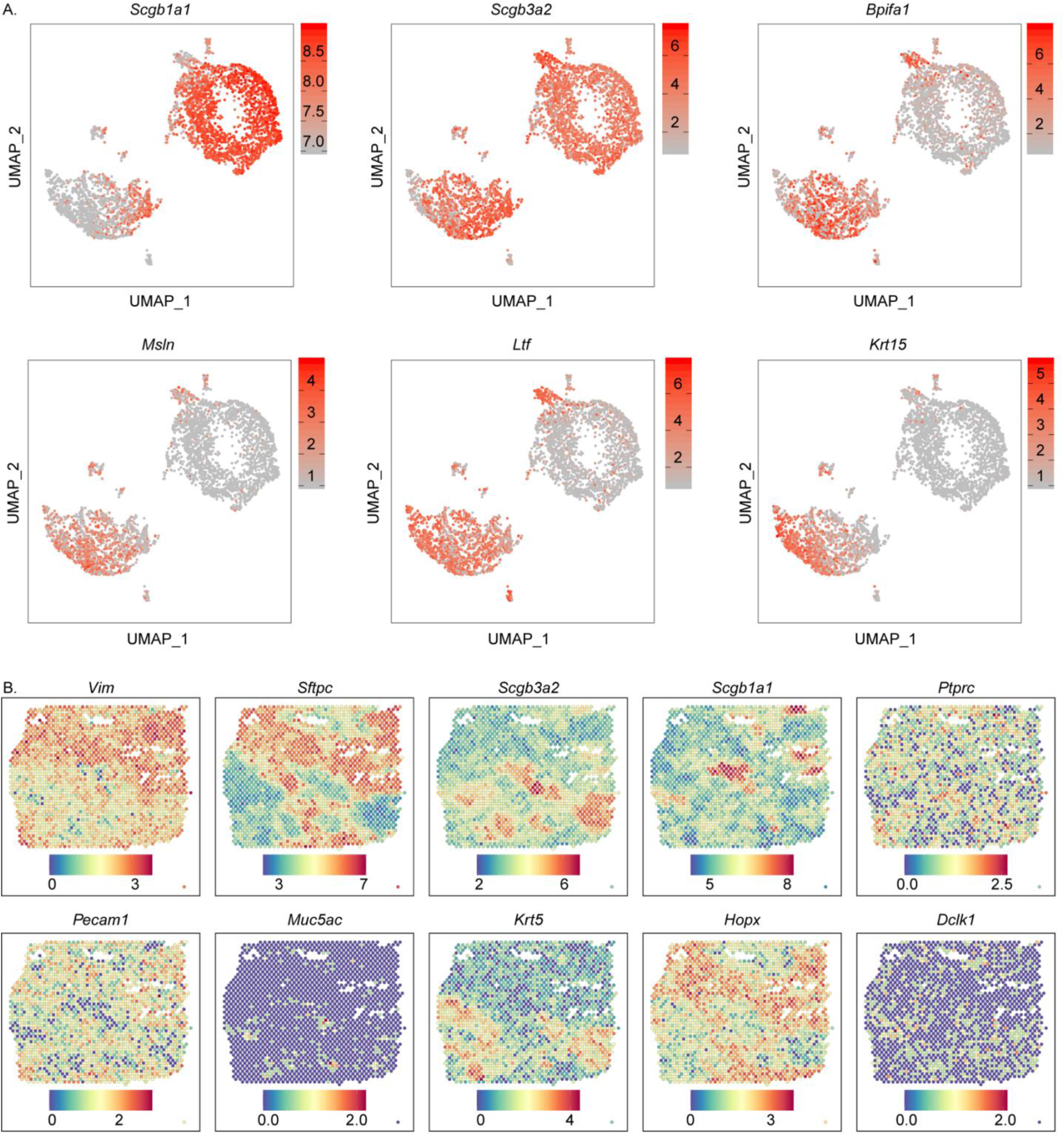
Assessment of transcriptional differences between club and IS populations. (A) Feature plot comparing expression of selected club and serous cell-specific genes between cell types. (B) Spatial feature plot comparing expression of selected cell-specific genes at 14 days following PR8 exposure: *Krt5* (basal), *Scgb3a2* (serous and club), *Bpifa1* (serous), *Muc5ac* (goblet), *Scgb1a1* (club), *Vim* (Fibroblast), *Pecam1* (endothelial), *Ptprc* (Immune).

**Supplemental Fig. 3.**
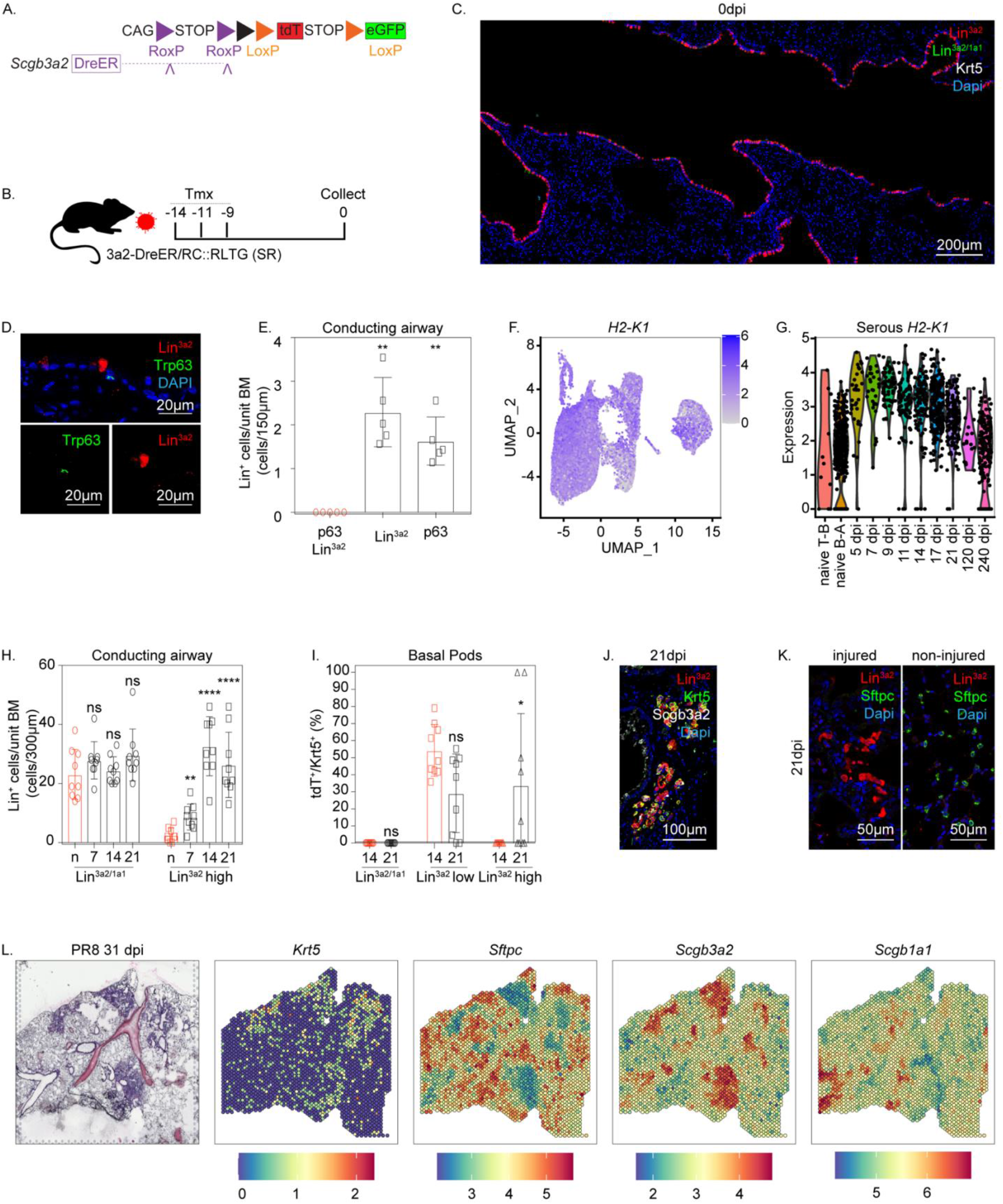
Further characterization of DR mice in the context of influenza induced lung injury. (A) Schematic illustration of recombinase driver and reporter allele used in Dre recombinase (SR) mice. (B) Experimental design. SR mice (n = 5 per experimental group) were treated with 3 doses of TM to assess degree of non-specific recombination. (C) Representative immunofluorescence localization of lineage reporters (green or red) with Krt5 (white) in SR mice at steady state. (D) Representative immunofluorescence colocalization of tdT (Lin^3a2^; red) with Trp63 (green) within conducting airway epithelium at steady state. (E) Contribution of lineage-tagged populations to Trp63+ progenitors within conducting airway epithelium at steady state. (F) Feature plot showing H2-K1 expression between cell types. (G) Violin Plot comparing expression of H2-K1 within subsetted serous cells at indicated time points after PR8 infection. (H) Contribution of lineage-tagged populations to hBC in airways as a function of time after PR8 infection (G). N=3 per condition, with significance determined by Mann-Whitney U-test. (I) Contribution of lineage-tagged populations to hBC in alveolar ‘pods’ as a function of time after PR8 infection (G). N=3 per condition, with significance determined by Mann-Whitney U-test. (J) Representative immunofluorescence colocalization of tdT (Lin^3a2^; red) with Krt5 (green) & Scgb3a2 (white) among lineage-positive alveolar clusters 21 days after PR8 infection. (K) Representative immunofluorescence colocalization of tdT (Lin^3a2^; red) with Sftpc (green) among lineage-positive alveolar clusters 21 days after PR8 infection. (L) Spatial gene expression and corresponding immunofluorescence of cell type-specific markers 31 days following exposure to PR8: *Krt5* (basal), *Scgb3a2* (serous and club), *Scgb1a1* (club) and *Sftpc* (AT2).

**Supplementary Fig. 4.**
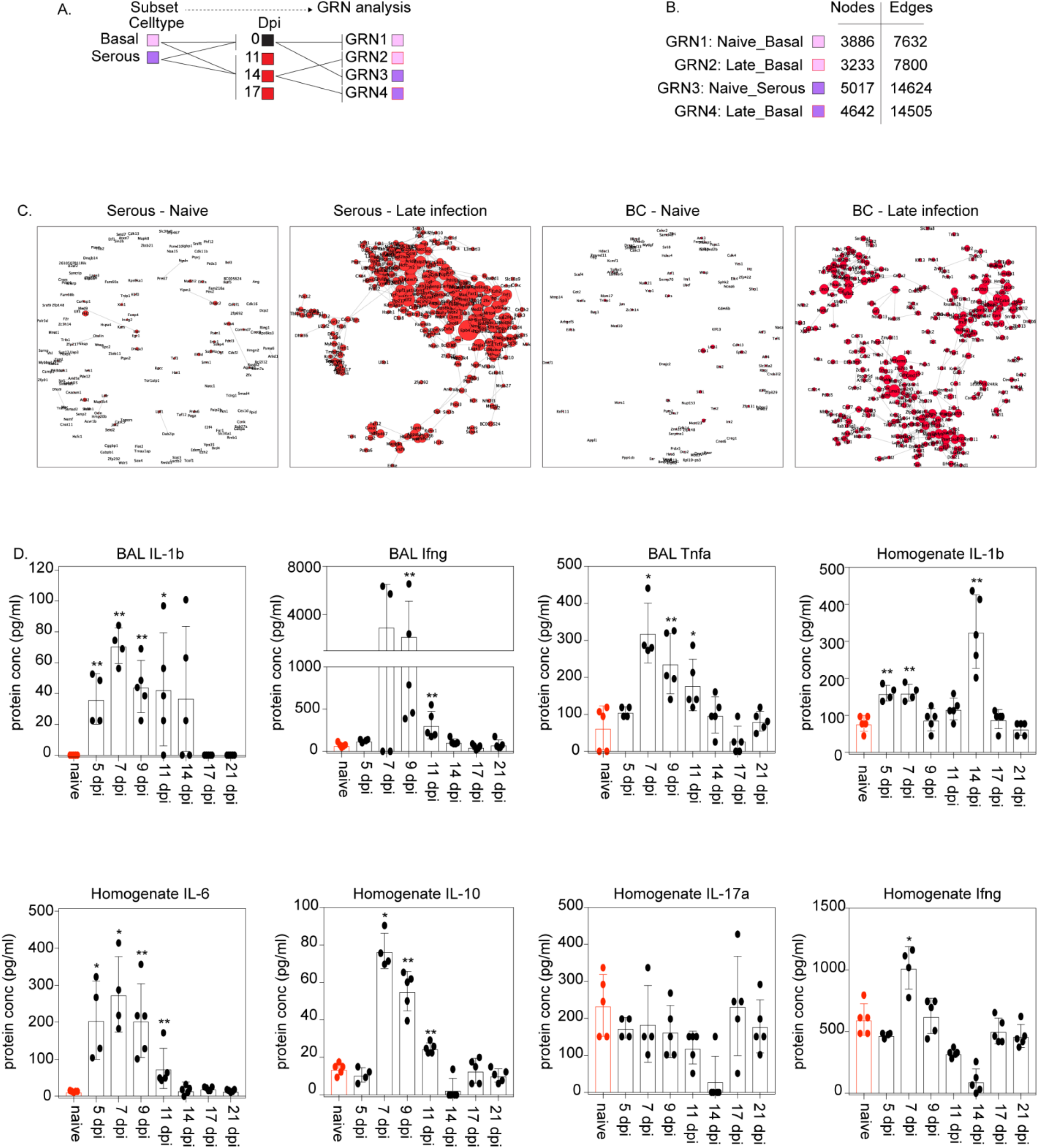
Gene regulatory network analysis of PR8 infected BC reveal increased immunoreactivity following injury. (A) Experimental design for generation of GRNs. ScRNAseq data were first subsetted by cell type followed by segregation into early vs late recovery time points, resulting in the creation of four networks for downstream analysis. 392 Serous and 513 BC were input for early timepoints; 412 Serous and 462 BC were input for late timepoints. (B) Table showing node and edge number after homogenization of late time points to their respective naïve controls. (C) Visualization of GRNs. Node and font sizes reflect degree centrality and strength of connectivity. The top five genes with the highest delta centrality value relative to naïve controls are highlighted in green, with remaining genes shown in red. (D) Assessment of cytokine levels in BALF & lung homogenate during recovery from PR8 infection. 4-5 biological replicates were used per timepoint with statistical significance determined by Mann-Whitney U-test.

**Supplemental Fig. 5.**
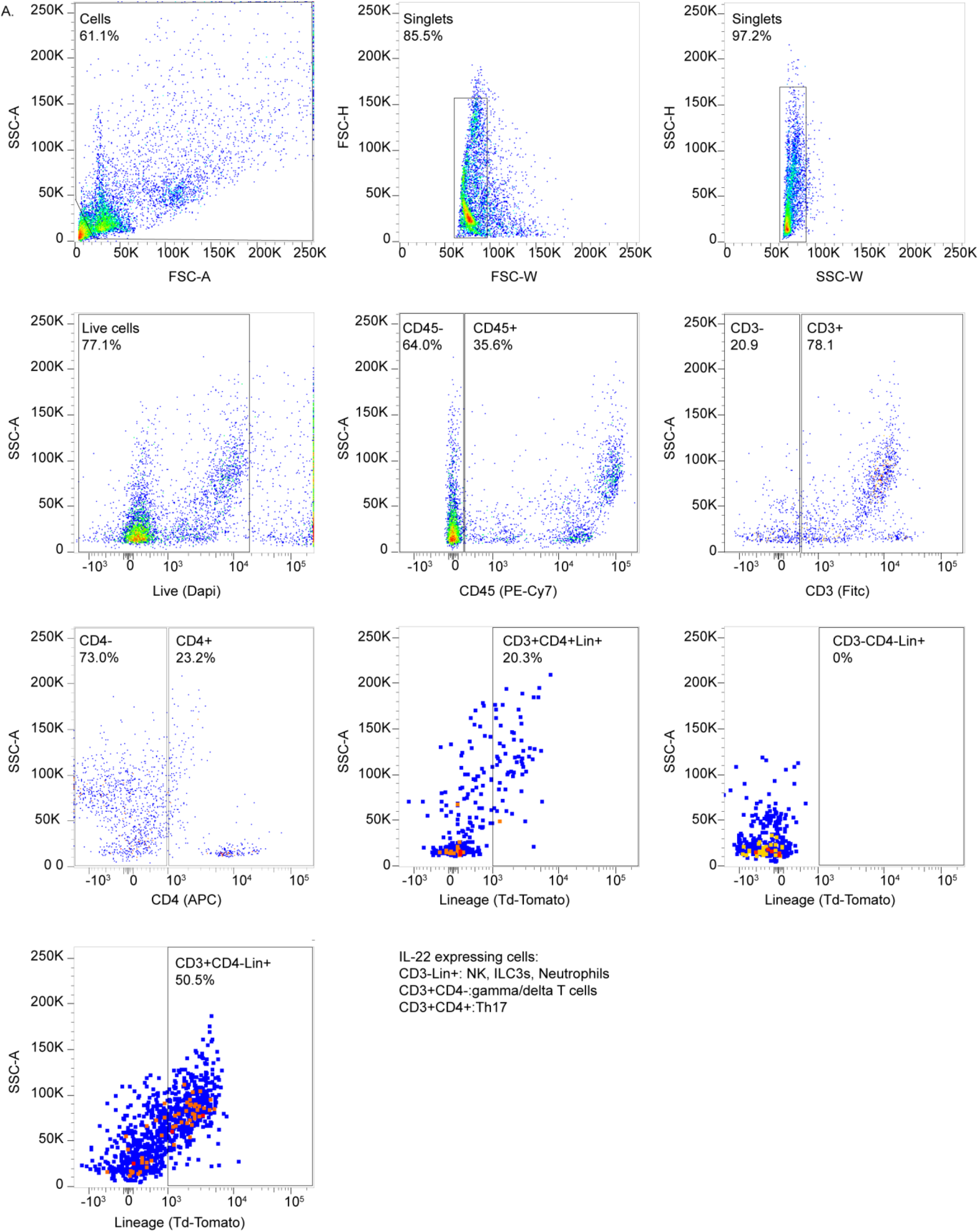
Gating strategy for quantification of IL-22 expressing subsets using flow cytometry. (A) Gating strategy to assess different IL-22 expressing cell types via flow cytometry. Cells were subsetted into CD3^-^Lin^+^, CD3^+^CD4^-^Lin^+^ and CD3^+^CD4^+^Lin^+^ populations to delineate non-T lymphoid, γδT and Th17 cells respectively.

**Supplemental Fig. 6.**
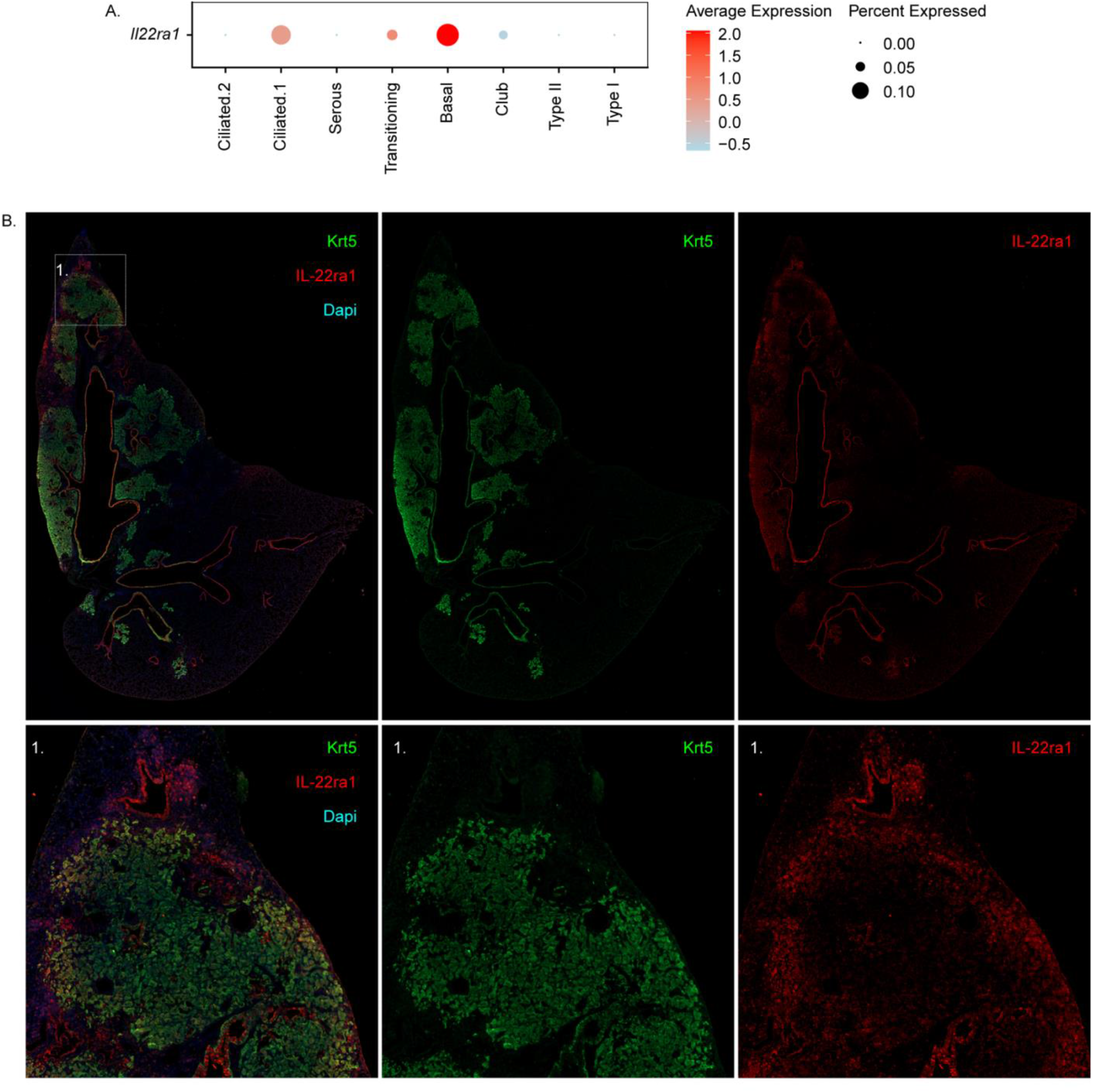
Visualization of IL-22ra1 positive and negative BC regions during repair. (A) Dot plot comparing expression of IL-22ra1 between epithelial subsets. (B) Representative immunofluorescent colocalization of IL-22ra1 (red) and Krt5 (green) in BC-rich alveolar region of PR8-infected mouse lung.

**Supplemental Fig. 7.**
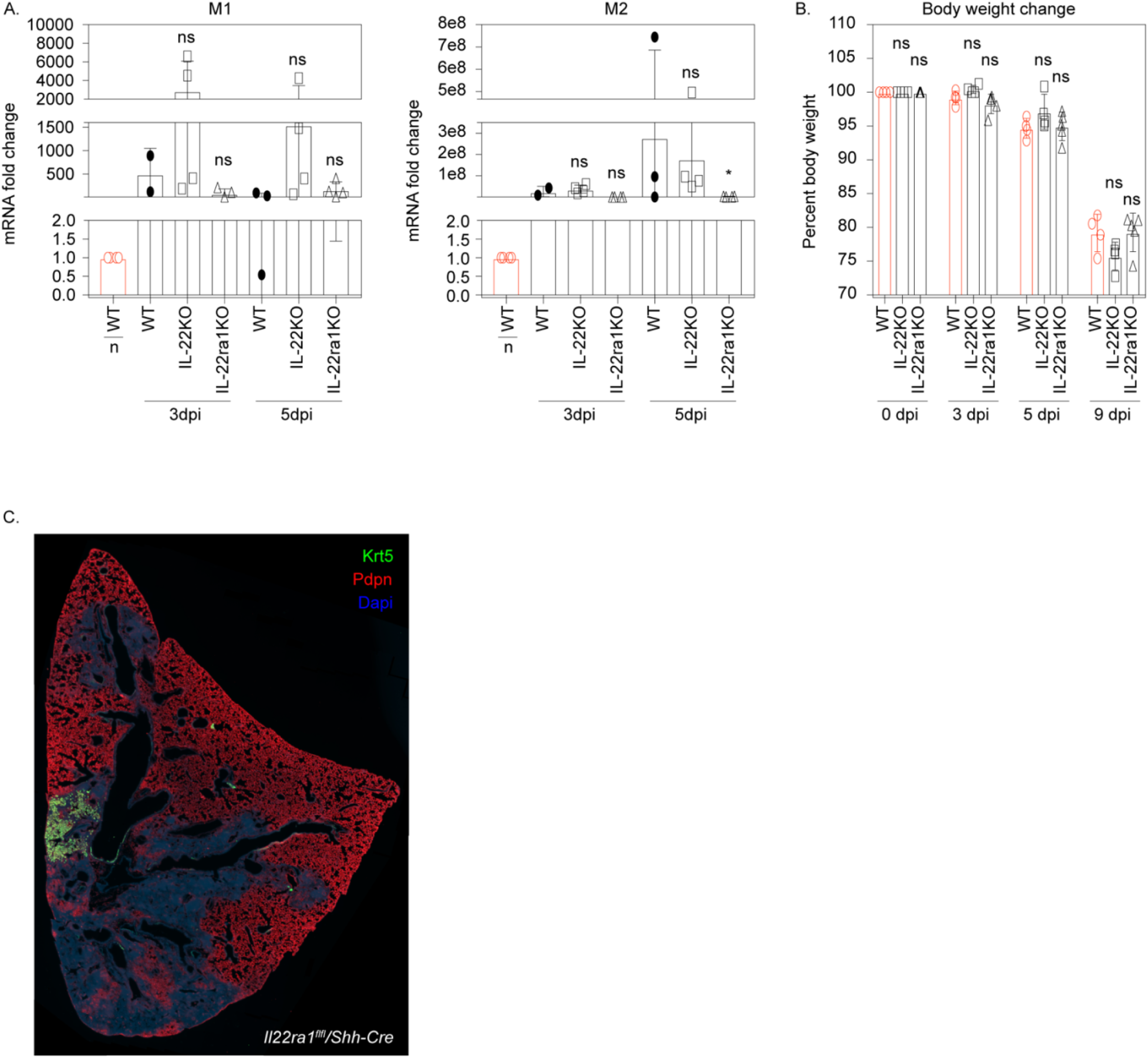
Additional phenotyping for IL-22 LOF & IL-22r cLOF. (A) qPCR detection of viral gene mRNA in total lung RNA of PR8-infected WT, IL-22 LOF, and IL-22r cLOF mice. 3-5 biological replicates were used per condition. Statistical analysis was performed by Mann-Whitney U-test. (B) Body weight changes of WT, IL-22 LOF, and IL-22r cLOF mice at different points during influenza induced acute lung injury. (C) Representative immunofluorescence localization of Pdpn (red) and Krt5 (green) in lungs of IL-22r cLOF mice 14 days post-PR8 infection.

## RESOURCE AVAILABILTY

### Lead Contact

Barry R. Stripp, Ph.D.

Cedars-Sinai Medical Center 8700 Beverly Blvd. AHSP Rm A9401 Los Angeles, CA 90048

### Materials availability

All unique/stable reagents generated in this study are available from the Lead Contact with a completed Materials Transfer Agreement.

### Data and Code availability

The data discussed in this publication have been deposited in NCBI’s Gene Expression Omnibus (Edgar et al., 2002) and are accessible through GEO Series accession number GSE184384 (https://www.ncbi.nlm.nih.gov/geo/query/acc.cgi?acc=GSE184384).

URL link to Github for code: https://github.com/BeppuAN/20220912_GEO-asssession-GSE184384

## EXPERIMENTAL MODEL AND SUBJECT DETAILS

### Mouse strains

Mice aged 8-12 weeks were used according to Institutional Animal Care and Use Committee-approved protocols. *Sftpc-CreER* (Rock et al., 2011), *Scgb1a1-CreER* (Rawlins et al., 2009),*Krt5-CreER* (Van Keymeulen et al., 2011), *IL-22*^*Cre*^ mice (Ahlfors et al., 2014), and *Shh-Cre* (Harfe et al., 2004) were obtained from Jackson Labs. *IL-22ra1*^*fl/fl*^ (Zheng et al., 2016) were provided by Jay Kolls. These mice were bred to either *ROSA-26-mTmG* (Muzumdar et al., 2007). *ROSA-26-tdTomato* (supplied by D. Jiang and P.W. Noble) or *RC::RLTG* (Plummer et al., 2015) mice for lineage-tracing experiments. *Scgb3a2-DreER* mice were generated by Jackson Labs using a CRISPR/Cas9 knock-in strategy. Mice were produced by inserting an IRES-DreERT2 construct into the 3’ UTR of the mouse *Scgb3a2* gene resulting in TM inducible DRE recombinase activity under the control of the *Scgb3a2* promoter. 4 Founders were identified following embryotic manipulation that carried the desired genetic permutation which was confirmed though long-range qPCR. Founders were back crossed to Bl6 mice to generate N1 progeny which were then bred to mice containing at least one copy of both *Scgb1a1-CreER* and *RC::RLTG* to generate the first generation of experimental mice for in vivo fate mapping experiments. For all experiments, 8-12 week-old mice were used and both male and female were included.

## METHOD DETAILS

### Tamoxifen

Tamoxifen (TM) was dissolved by sonication at a concentration of 20mg/ml in corn oil until clear and stored at -80°C in 50ml aliquots. A washout of 9 days following the last treatment with TM was used for initial lineage tracing analysis prior to infection. For these experiments, DR mice were injected intraperitoneally using a 27 gauge needle 3 times at a dose of 0.225mg/g mouse body weight. Additional experiments were performed on DR mice with extended TM washout, to confirm that lineage tracing experiments were not confounded by residual TM and associated nuclear Cre recombinase activity following exposure to virus. For these experiments, TM washout period was extended to 28 days and mice were treated by oral gavage at a dose of 0.225mg/g body weight. No differences in lineage tracing outcome were observed between animals undergoing 9 day vs 28 day TM washout prior to PR8 infection.

### Influenza Virus inoculation

A mouse-adjusted variant of the 1918 H1N1 influenza (Puerto Rico 8; PR8) was used to infect mice and induce acute lung injury. A single treatment of 100PFU/50μl was administered intratracheally into mice placed into a surgical plane through inhalation exposure to isoflurane. Mouse weight was monitored over a 14-day period to verify uniformity of infection between individuals. Expression of viral M1 & M2 gene transcripts were assessed from lung homogenates of representative mice to further confirm that a reproducible level of infection has occurred under all experimental conditions and between mouse genotypes. Mouse lung biopsies were collected at time points that correspond with hyperplasia of basal-like cells in distal lung tissue of PR8-infected mice.

### Preparation of single cell suspensions for Single cell RNAseq

#### Type II & immune subsets

Mouse lung biopsies were collected at the indicated time points: naïve, 3, 5, 7, 9, 11, 14, 17, 21, 60, 120 and 240 days post infection. A sample size of 5 C57/Bl6 WT mice were used for each timepoint. Cell suspensions from each condition were pooled together prior to cell sorting. On the day of biopsy collection, the entire mouse lung was separated from the chest cavity and stored in a conical containing 4° C 1XHBSS. Isolated lung lobes were intratracheally instilled with a 3ml mixture containing 1U elastase/1ml 1XHBSS for 30 minutes at 37° C. The crude cell suspension underwent mechanical agitation prior to incubation in dissociation solution with a final composition of 1X Liberase/1X HBSS for 30 minutes at 37°C. Dissociation buffer was quenched with a solution containing 2% FBS/1mM EDTA/1X HBSS on ice. Cells were filtered through a 70μm nylon mesh to remove undigested tissue. Cells were centrifuged at 500xg for 10 minutes and resuspended in 1ml blood cell lysis solution for 1 minute to remove red blood cells from the suspension. Red blood cell lysis buffer was quenched using 25 ml 2% FBS/1mM EDTA/1X HBSS, followed by centrifugation at 500 x g for 10 minutes to pellet intact cells. When isolating epithelium, cells were resuspended in 1ml 2% FBS/1mM EDTA/1X HBSS and magnetic bed separation was performed to deplete CD31 and CD45 subsets. Cell suspensions were enriched for epithelial and immune populations by FACS, with selection for CD326^+^CD45^-^CD31^-^ and CD326^-^CD45^+^CD31^-^ to yield enriched epithelial and immune cell fractions, respectively. A minimum of 50,000 epithelial cells and 100,000 immune cells were sorted for each timepoint.

#### Conducting airway subsets

Single-cell RNA seq data was generated from cell suspensions enriched for conducting airway epithelium. The aforementioned cell suspension prep was performed on TM inoculated *Sftpc-CreER*/*ROSA-mTmG* and a gating strategy was used to enrich for tdT negative epithelium (i.e. CD326^+^CD45^-^CD31^-^tdT^-^) by FACS. For these experiments, a sample size of 3 *Sftpc-CreER*/*ROSA-mTmG* were pooled together prior to cell sorting. Cells were collected at the same indicated time points (naïve, 3, 5, 7, 9, 11, 14, 17, 21, 60, 120 and 240 days post infection) and a minimum of 50,000 cells were collected per timepoint. In experiments where isolation of conducting airway epithelium was not possible using *Sftpc-CreER*/*ROSA-mTmG* (Fig. 7H) an alternative gating strategy was used to deplete for Type II cells. A combination of Sca-1 and CD24 antibodies were used in lieu of the tdT lineage tag (i.e CD326^+^CD45^-^CD31^-^Sca1^+^CD24^+^).

### Bioinformatics of scRNAseq

#### Dataset Generation

Isolated single cell suspension was resuspended in 0.04% BSA at a concentration of 600cells/ul and were used to generate GEMS using a 10x genomics chromium controller. TotalSeq Hashtag Antibodies (Biolegend:15580#) were used to multiplex samples from isolated conducting airway epithelium (i.e. those cells isolated from TM-inoculated *Sftpc-CreER*/*ROSA-mTmG* mice described in the previous section). ScRNAseq datasets described in Fig. 1A and Fig. 4C were generated using Chromium Single Cell 3’ v2 chemistry. ScRNAseq dataset described in Fig. 7H were generated using Chromium Single Cell 3’ v3 chemistry. Libraries were generated as per manufactures instructions. Sequencing was performed on a Novaseq 6000 using a S4 150 and 28 + 90 bp paired-end setting. Alignment was performed using ‘Cellranger count’ function provided in 10x Genomics single cell gene expression software.

#### Secondary analysis

The R package ‘Seurat’ (Butler et al., 2018; Stuart et al., 2019) was used to apply standard quality control metrics and unsupervised clustering for generation of initial UMAP projections. The R package ‘FGSEA’ (Korotkevich et al., 2021) was used to perform gene set enrichment analysis. Single cell datasets were subsetted based on cell type and changes in gene expression was assessed at different points during flu infection using the hallmark gene sets from the molecular signature database (Liberzon et al., 2015; Subramanian et al., 2005). The R package ‘Velocyto’ and ‘scVelo’ (Bergen et al., 2020; La Manno et al., 2018) was used to perform the single cell trajectory analysis. Fastq files were aligned to a separate reference transcriptome containing both splice and unspliced gene variants. Before calculating RNA velocity, cell types of interest were subsetted from the compiled epithelial dataset and further subsetted based on timepoint. This was primarily done to reduce the computation required by the Velocyto package. Degree node centralities were calculated from gene regulatory networks generated by the package ‘Bigscale2’ (Iacono et al., 2018). The ‘Bigscale2’ package has functionality to calculate difference in node centralities between two conditions and this was used to generate ranked lists of the top 300 delta degree node centralities between early and late infection. The resultant rank list was used to assess enriched GO terms using a PANTHER based over-representation test produced by the gene ontology consortium (Ashburner et al., 2000; Gene Ontology, 2021; Mi et al., 2013). Gene networks were visualized using Cytoscape_v3.8.1.

#### Spatial transcriptomics

Frozen 10μm sections from 14 days post PR8 infected mouse lungs were placed within the frames of the capture areas on the active surface of the Visium spatial slide. Tissue sections were fixed in methanol and stained with H&E. Bright-field images of stained sections in the fiducial frames were collected in 40x fields using Zeiss Axioscan Z1 microscopy. Stained tissue sections were permeabilized for 30 minutes and mRNA was released to bind oligonucleotides on the capture areas. Single cell RNA-seq libraries were prepared as per manufacture instructions and sequenced on a Novaseq 6000 using SP 28 + 90 bp paired-end reads. Count matrices were generated using the ‘spaceranger count’ function in Space Ranger 1.0.0. The resulting data were processed in Seurat. Mouse scRNASeq clusters described in Fig.1A and Fig. 4C were transferred to spatial transcriptomic sample. The transfer function generates a probability score for each spot and its association with a given scRNASeq cluster. The spot is assigned to the cluster with the highest score and mapped back to the spatial transcriptomic sample image.

### Immunofluorescent microscopy

#### Immunofluorescence staining

To prepare mouse lung for histology, we inflation fixed freshly dissected mouse lung through instillation of 1ml 4% paraformaldehyde directly into a cannulated mouse trachea. After incubating the lungs for 24 hours, the left lobe from each mouse was separated into a labeled cassette and stored in either 1X PBS for immediate tissue processing or in 70% EtOH for long term storage. Tissues were dehydrated using the ASP300 tissue processor(Leica). After processing, left lobes were embedded in paraffin wax and sectioned at 7-9μm thickness using a HM 325 rotary microtome(Leica). Sectioned slides were dried at room temperature until needed. After selection of an appropriate panel, tissue sections were deparaffinized using the Shandon veristain Gemini ES (Thermofisher, cat:A7800013). Slides were transferred into a reservoir containing either citrate or tris-based antigen retrieval solution(vector labs, cat:3300/3301) and heated using a pressure cooker (Biovendor, cat:RR2100-EU). The slides were then blocked with 2% BSA for 30 minutes. Tissue sections were incubated in primary antibody overnight at 4°C. Cells were then washed 5 times with 1X PBS and incubated in solution containing both alexaflour conjugated secondary antibody and DAPI for 2 hours. Slides were cover-slipped using Fluromount-G (EMS, cat:7984-25). Images were taken using either Zeiss 780 confocal microscope or the Zeiss Axioobserver Z1 inverted microscope.

#### Following are primary antibodies used

Chicken polyclonal anti-eGFP (1:1000, Abcam, Ab13970); Chicken Polyclonal anti-Keratin 5 (1:500, BioLegend, 905901); Mouse monoclonal anti-eGFP AF488 conjugated (1:500, Santa Cruz, Sc-9996); Rat monoclonal anti-IL-22ra1 (1:200, R&D systems, MAB42341); Goat polyclonal anti-tdTomato (1:500, Sicgen, Ab8181-200); Goat polyclonal anti-p63(1:500, Santa Cruz, Sc-8609); Goat Polyclonal UGRP1/SCGB3A2 (1:1000, R&D Systems, AF3465); Syrian hamster Monoclonal anti-Pdpn (1:1000, LifeSpan Biosciences, LS-C143022-100); Rabbit Polyclonal anti-SCGB1A1 (1:500, Proteintech, 10490-1-AP); Rabbit Polyclonal anti-tdT(1:500, Rockland, 600-401-379); Rabbit Polyclonal anti-Msln (1:500, Thermo Fisher Scientific, PA5-79698); Rabbit Polyclonal anti-Ltf (1:200, Thermo Fisher Scientific, PA5-95513); Rabbit Polyclonal anti-Bpifa1 (1:200, Sigma-Aldrich, AV42475); Rabbit polyclonal anti-Keratin 5 (1:500, Cell Marque, EP1601Y); Rabbit polyclonal anti-Keratin 5 (1:500, Santa Cruz, Sc-66856); Rabbit polyclonal anti IL-22 (1:200, Abcam, ab18499); Rabbit polyclonal anti-Ki67 (1:1000, Ebioscience, 14-5698-82). Following are secondary antibodies used: Goat anti-Chicken Alexa Fluor 488(1:500, Thermo Fisher Scientific, 6100-30); Goat anti-Hamster Alexa Fluor 488 (1:500, Thermo Fisher Scientific, A-21110); Donkey anti-Rabbit Alexa Fluor 488 (1:500, Thermo Fisher Scientific, A-21206); Donkey anti-Goat Alexa Fluor 555(1:500, Thermo Fisher Scientific, A-21432); Donkey anti-Rabbit Alexa Fluor 555 (1:500, Thermo Fisher Scientific, A-31572); Goat anti-Chicken Alexa Fluor 568 (1:500, Thermo Fisher Scientific, A-11041); Donkey anti-Rat Alexa Fluor 594(1:500, Thermo Fisher Scientific, A-21209); Goat anti-Hamster Alexa Fluor 594 (1:500, Thermo Fisher Scientific, A-21113); Goat anti-Chicken Alexa Fluor 647 (1:500, Thermo Fisher Scientific, A-21449); Donkey anti-Rabbit Alexa Fluor 647 (1:500, Thermo Fisher Scientific, A-31573); Donkey anti-Goat Alexa Fluor 647 (1:500, Thermo Fisher Scientific, A-31573); Donkey anti-Goat Alexa Fluor 647 (1:500, Thermo Fisher Scientific, A-21447).

### Lineage tracing analysis

#### Quantification of Lin^3a2^ area as a percentage of Krt5^+^ area

The composite image of lung sections stained with Rabbit anti-Krt5(green; Cell Marque: EP1601Y), Gt anti-tdT(red; Scigen: Ab8181-200) and DAPI (blue) were imported into Fiji image analysis software as separate images based off their respective channels. Positive staining for Krt5 and tdT was determined using Fiji’s ‘Threshold’ function: Image -> adjust -> threshold. A separate image showing overlapping pixels between the Krt5 and tdT image was generated using the ‘AND’ operation in Fiji’s image calculator function: Process -> Image calculator. Areas containing negatively stained nuclei (or ‘holes’) were resolved using Fiji’s Watershed function: Process-> Binary-> Watershed. Krt5 and Krt5-tdT overlapping areas were measured using the following sequence of features: Analyze -> Analyze particles. Percent tdT/Krt5 area was calculated by Krt5-tdT area as a function of total Krt5 area.

#### Quantification of cells per unit BM

To score the number of cells per unit BM, 20 lines along the basement membrane was measured using the ‘segmented line tool’ in the Fiji image analysis software(Schindelin et al., 2012) and recorded into a google spreadsheet. The number of lineage tagged tdT and eGFP from DR mice was counted along a basement membrane length of 300μm.

**Quantification of Pod region percent totals:** Pods were defined as at least five continuous Krt5/eGFP/tdT-immunoreactive cells in alveolar regions that were not associated with pre-existing bronchial epithelium. 100 pod cells per region of interest were counted using Fiji’s ‘multi-point’ function and recorded into a google spreadsheet. Then the percentage of either tdT low, tdT high or eGFP positive cells were assessed based on the function of counted pod cells.

### Quantification of Krt5 area in *IL-22*^*Cre*^ and *IL-22ra1*^*fl/fl*^/*Shh-Cre*

The composite image of lung sections stained with Ck anti-Krt5(green; BioLegend: 905901), Hm anti-Pdpn(red; LifeSpan Biosciences: LS-C143022-100) and DAPI (blue) were imported into Fiji image analysis software as separate images based off their respective channels. Images were converted into greyscale using the following sequence of image processing features: Image -> Type -> 8bit.

Area’s containing positive stain were then converted into binary black/white images using the following sequence of images processing features: Process -> Binary-> Make Binary. Area of each channel was measured and recorded into a google spreadsheet using the following sequence of features: Analyze -> Analyze particles. Quantification of Krt5 area was calculated by measuring the area of Krt5^+^ immunoreactive pods in proportion to the area of damaged regions demarcated by DAPI^+^PDPN^-^ stain within alveolar epithelium using Fiji software.

### Quantitative real-time PCR

RNA was extracted from homogenized snap frozen superior lobes of either *IL-22*^*Cre*^ homozygous, *IL-22ra1*^*fl/fl*^/*Shh-Cre* or WT controls using RNAeasy mini kit (Qiagen, cat: 74106). Isolated RNA was converted into cDNA using a iScript cDNA synthesis kit (Bio-Rad, cat:1708891). Gene expression analysis was performed using SYBR Green PCR master Mix (Thermo Fisher Scientific, cat: 4309155) and analyzed on the 7500 fast Real-Time PCR system (Thermo Fisher Scientific).

### Flow cytometry of IL-22 expressing cells

*IL-22*^*Cre*^ homozygous mice were purchased and bred to heterozygosity with another strain expressing at least one copy of *ROSA-tdT* to generate *IL-22*^*Cre*^/*ROSA-tdT* mice. Mouse lungs were collected from steady state and from mice infected with influenza for 14 days. Lungs was collected and processed into single cell suspension as described above in the ‘Preparation of single cell suspensions for Single cell RNAseq’ section. An immune panel capable of delineating between T helper and non-T-helper subsets (CD3^+^CD4^+^tdT^+^ and CD3^+^CD4^-^ tdT^+^ respectively) was used to determine the main IL-22 expressing cells using flow cytometry.

### Multiplex protein assays

Mouse bronchial alveolar lavage and mouse left lobes were collected at the following time points: naïve, 3, 5, 7, 9, 11, 14, 17, 21dpi. BAL was prepared through instillation of intubated mouse trachea with 1ml 1X dPBS three times. Instilled 1X dPBS was transferred into a 1.5 microfuge tube. To prepare lung homogenates, dissected lobes were transferred into collection tubes containing 1.4 mm ceramic beads (Lysing matrix D, MP Biomedicals Cat: 116913100) and homogenized using mechanical agitation (MP Benchtop Homogenizer, MP biomedicals, Cat: 6VFV9). Samples were centrifuged at 600xg to pellet cells. Supernatants were transferred into a separate 1.5mL collection tube. A multiplexed protein assay (Bio-plex Pro, Bio-Rad, Cat: 171304070, M69999997NY) was performed to assess changes in expression of the following cytokines: IL-1b, IL-6, IL-10, IL-17, IL-22, Tnfα and Ifnγ. Samples were processed for analysis as per manufactures instructions. Cytokine levels were quantified using the fluid flow-based microplate reader (Bio-plex 200, Luminex, Cat:171000201).

## QUANTIFICATION AND STATISTICAL ANALYSIS

Detailed descriptions relating to quantification and statistical analysis of each experiment is documented in the above Methods section. Statistical analysis from graphs generated by wet lab experiments was performed in Graphpad Prism 7. Variance in datapoints between conditions is represented by Mean +/-SEM. Statistical analysis for single cell RNA seq data was performed using pipelines for statistical analysis inherit to each package.

## ADDITIONAL RESOURCES

### KEY RESOURCES TABLE

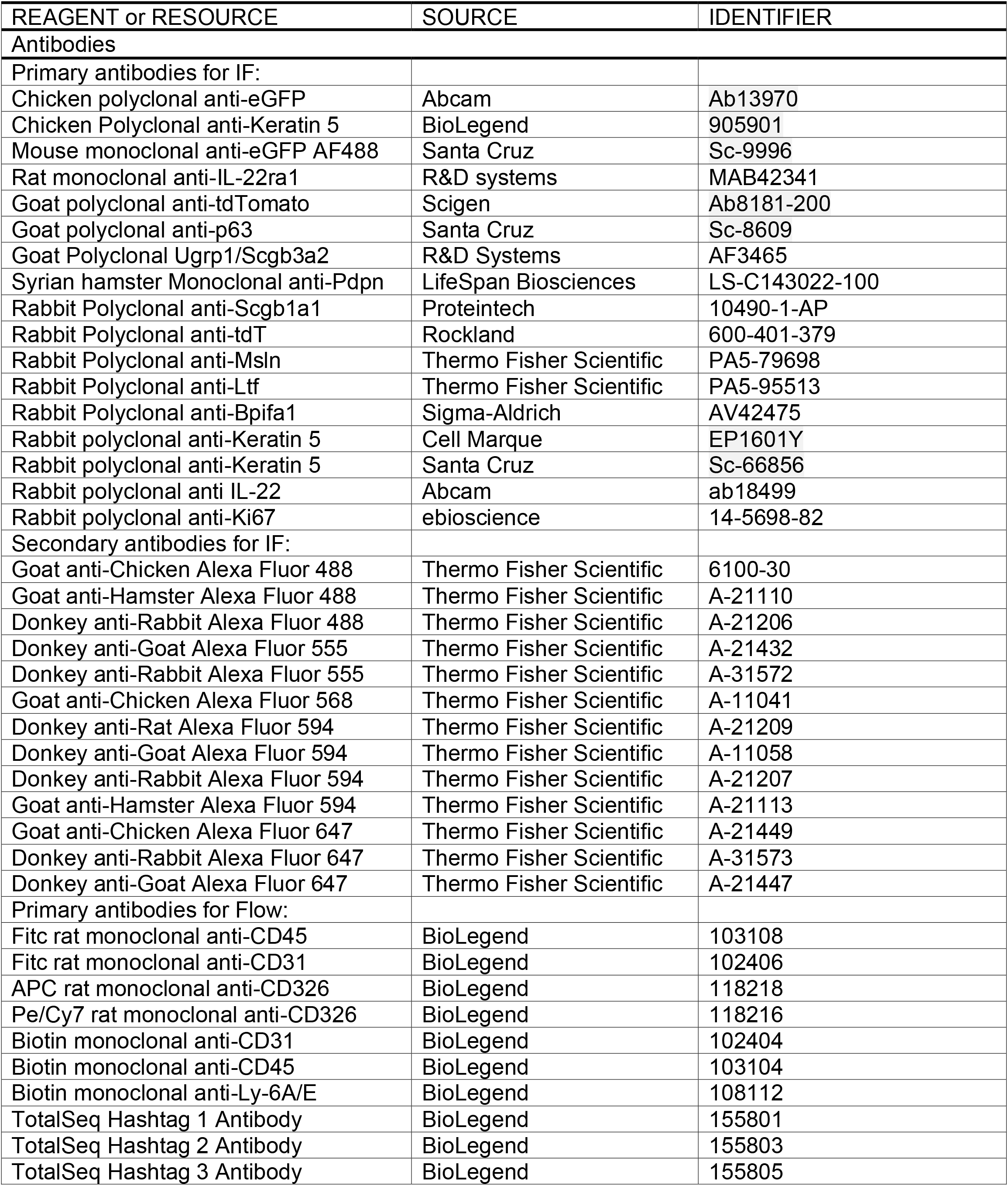
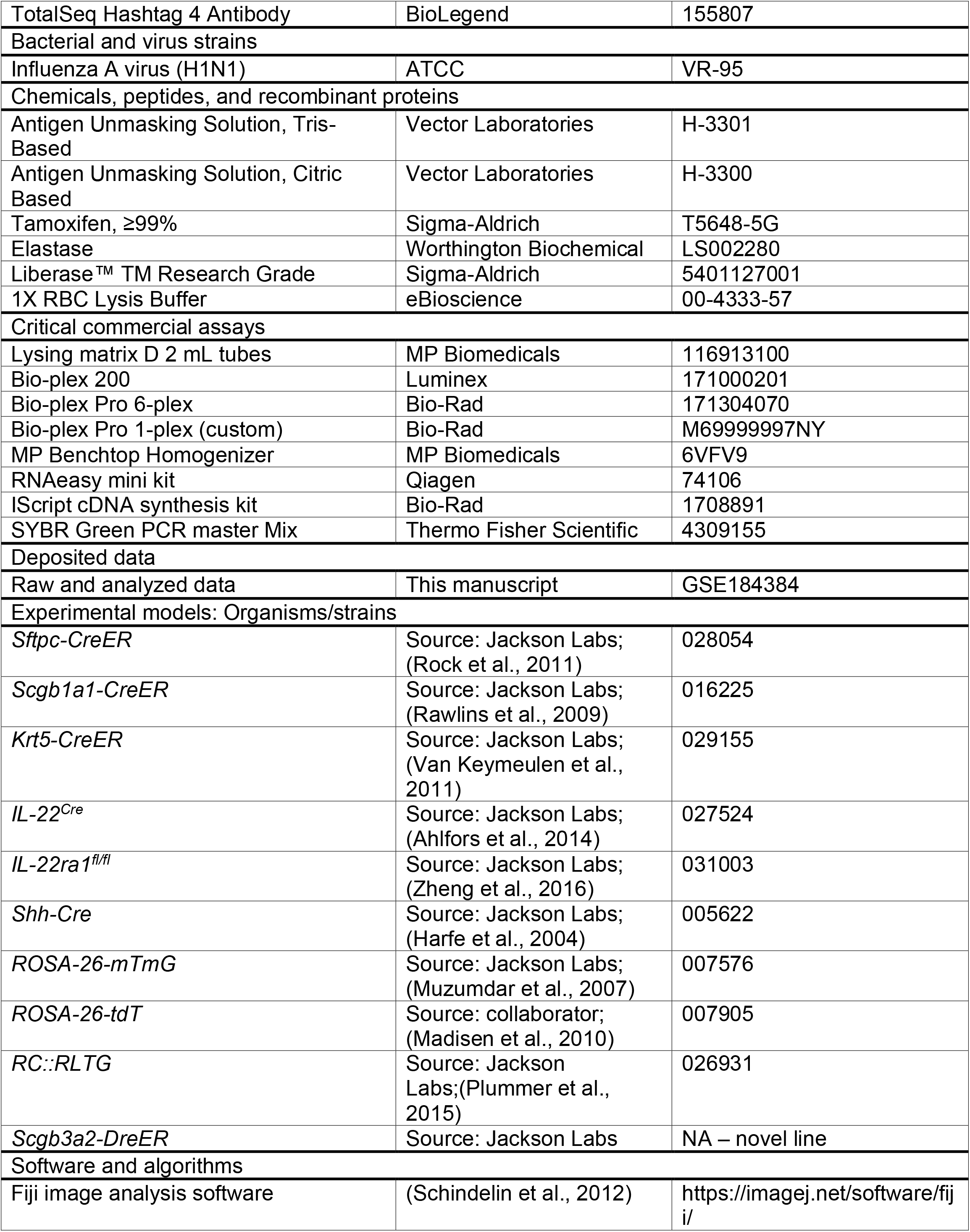
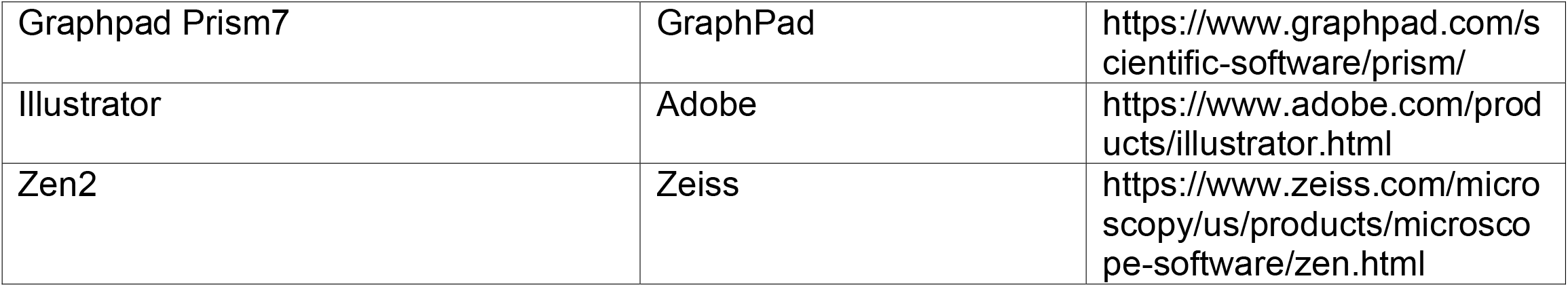

